# Functional characterization of the biogenic amine transporter system on human macrophages

**DOI:** 10.1101/2021.09.08.459459

**Authors:** Phillip M Mackie, Adithya Gopinath, Dominic M Montas, Alyssa Nielsen, Rachel Nolan, Kaitlyn Runner, Stephanie Matt, John McNamee, Joshua Riklan, Kengo Adachi, Andria Doty, Adolfo Ramirez-Zamora, Long Yan, Peter J Gaskill, Wolfgang J Streit, Michael S Okun, Habibeh Khoshbouei

**Affiliations:** Department of Neuroscience, University of Florida College of Medicine, Gainesville FL, 32610; Department of Pharmacology and Physiology, Drexel University College of Medicine, Philadelphia PA, 19102; Neuronal Signal Transduction Group, Max Planck Florida Institute for Neuroscience, Jupiter FL 33458; Interdiscipinary Center for Biotechnology Research, University of Florida, Gainesville FL, 32610; Department of Neurology, University of Florida College of Medicine, Gainesville FL, 32610; Norman Fixel Institute for Neurological Disease, Department of Neurology, University of Florida College of Medicine, Gainesville FL, 32610; Center for Translational Research in Neurodegenerative Disease, University of Florida College of Medicine, Gainesville FL, 32610

**Author notes:** To whom correspondence should be addressed Habibeh Khoshbouei, Phillip M Mackie.

## Abstract

Monocyte-derived macrophages are key players in tissue homeostasis and disease regulated by a variety of signaling molecules. Recent literature has highlighted the ability for biogenic amines to regulate macrophage functions, but the mechanisms governing biogenic amine signaling on and around immune cells remains nebulous. In the central nervous system, biogenic amine transporters are regarded as the master regulators of neurotransmitter signaling. While we and others have shown macrophages express these transporters, relatively little is known of their function on these cells. To address these knowledge gaps, we interrogated the function of norepinephrine (NET) and dopamine (DAT) transporters on human monocyte-derived macrophages. We found that both NET and DAT are present and can uptake substrate from the extracellular space at baseline. Not only was DAT expressed in cultured macrophages, but it was also detected in a subset of intestinal macrophages *in situ.* Surprisingly, we discovered a NET-independent, DAT-mediated immuno-modulatory mechanism in response to lipopolysaccharide (LPS). LPS induced reverse transport of dopamine through DAT, engaging autocrine/paracrine signaling loop that regulated the macrophage response. Removing this signaling loop enhanced the pro-inflammatory response to LPS. Finally, we found that this DAT-immune axis was disrupted in disease. Collectively, our data introduce a novel role for DAT in the regulation of innate immunity during health and disease.

## Introduction

Monocytes and monocyte-derived macrophages (MDM) are a heterogenous population that serve as critical components of the immune system. Monocyte-derived macrophages arise when monocytes engraft into tissues to replenish the resident macrophage pool as in the gut, dermis, heart, and lung (reviewed in (Ginhoux and Guilliams 2016), or in response to inflammatory signals (Shi and Pamer 2011). Once engrafted, MDM become transcriptionally distinct from circulating monocytes adopting a unique phenotype depending on their microenvironmental niche (Lavin et al. 2015). Although MDMs in different tissues have specialized functions, their fundamental activities include phagocytosis, cytokine production and antigen presentation. Defective or aberrant macrophage functions have been associated with inflammatory (Na et al. 2019), autoimmune (Ma et al. 2019), and neurological diseases (Chiot et al. 2020, Jordão et al. 2019).

Macrophage functions are regulated by signaling molecules found in their local milieu. Classically, these factors are cytokines released from other immune cells, but macrophages can respond to a variety of other stimuli, including neurotransmitters released from peripheral nerve terminals (Sternberg 2006). In fact, the autonomic nervous system has long-been recognized for its ability to regulate the immune response (Borovikova et al. 2000, Elenkov et al. 2000). Biogenic amines, such as norepinephrine, in particular have been noted for their ability to dynamically regulate macrophage function (Scanzano and Cosentino 2015). Indeed, a subset of intestinal macrophages express beta-adrenergic receptors that engage a tissue-protective phenotype (Gabanyi et al. 2016). Additionally, adipose macrophages adjacent to sympathetic terminals express a functional norepinephrine transporter (NET) that modulates the pro-inflammatory state and thermogenesis (Pirzgalska et al. 2017), indicating that the biogenic amine transporter activity itself can influence the immune system.

Recently, dopamine has been reported to have its own immunomodulatory properties independent of norepinephrine (Torres-Rosas et al. 2014, Nickoloff-Bybel et al. 2019). Dopamine is found in the kidney, adrenal glands, carotid body, and gut in concentrations high enough to activate dopamine receptors (Matt and Gaskill 2019). Dopamine receptor activation has a wide range of effects on macrophage functions such as phagocytosis, cytokine profile, and inflammasome activation (Pinoli, Marino and Cosentino 2017, Nolan et al. 2019). The variance in the results might be partially explained by the different concentrations of dopamine used (and thus the activated dopamine receptor type) and whether the study was done in human primary immune cells, mice, or immortalized cell lines, or immune-like heterologous cells. The often-times conflicting results indicate that our knowledge of dopamine signaling on immune cells is incomplete.

In the CNS, dopamine signaling is a dynamic, tightly regulated process with the dopamine transporter (DAT) serving as the master regulator of dopamine transmission (Beckstead et al. 2004). DAT can regulate dopamine signaling by uptake of extracellular dopamine into the cell and efflux of intracellular dopamine out of the cell. Although macrophages express the dopamine transporter (DAT) and the proteins for synthesis and storage of dopamine (Gaskill et al. 2012, Mackie et al. 2018), how immune cells may regulate dopamine signaling via DAT, or how DAT activity itself may modulate macrophage phenotype is largely unknown. Such information on human primary macrophages would help elucidate a more comprehensive view of dopamine’s role in immune function and would be essential to help guide future studies targeting the dopamine system as an immunomodulator.

In this study, we aimed to first characterize the biology of biogenic amine transporters on primary human monocyte-derived macrophages. Employing multiple complementary approaches such as flow cytometry, qPCR, immunoblotting, fixed- and live-cell microscopy, we found that primary monocyte-derived macrophages from healthy human subjects express a functional NET and DAT, but not SERT. Furthermore, we found that, in addition to monocyte-derived macrophages *in vitro*, a subset of human intestinal macrophages that express DAT *in-situ*. While NET expression and its activity on human macrophages are known, the discovery of functional DAT on these cells was unexpected. Importantly, we discovered that DAT activity plays an integral part in modulating the macrophage response to endotoxin, independent of NET. We attributed this to LPS-induced DAT-mediated efflux of dopamine and enhancement of autocrine dopamine signaling. Finally, we found that this DAT-immune axis is perturbed in the context of Parkinson’s disease. Thus, our findings indicate the novel importance of DAT as a potential immunological rheostat.

## Results

### Human monocytes and macrophages express NET and DAT, but not SERT

In order to study the biology of biogenic amine transporters on primary human monocytes and monocyte-derived macrophages (MDM), we isolated peripheral blood mononuclear cells from peripheral blood (**Figure 1A**). Using a previously published protocol generated from our lab (Gopinath et al. 2020), we measured the percent of freshly isolated monocytes expressing either norepinephrine (NET) or dopamine (DAT) transporters. Approximately 18% of monocytes were DAT+; whereas approximately 4-5% of monocytes were NET+ and no monocytes were positive for serotonin transporter (SERT, **Figure 1B**, gating strategy in **Supplemental Figure 1**). Notably, these data imply the existence of a population of DAT+/NET- monocytes, suggesting DAT may have a role in monocyte function independent of NET.

**Figure 1:**
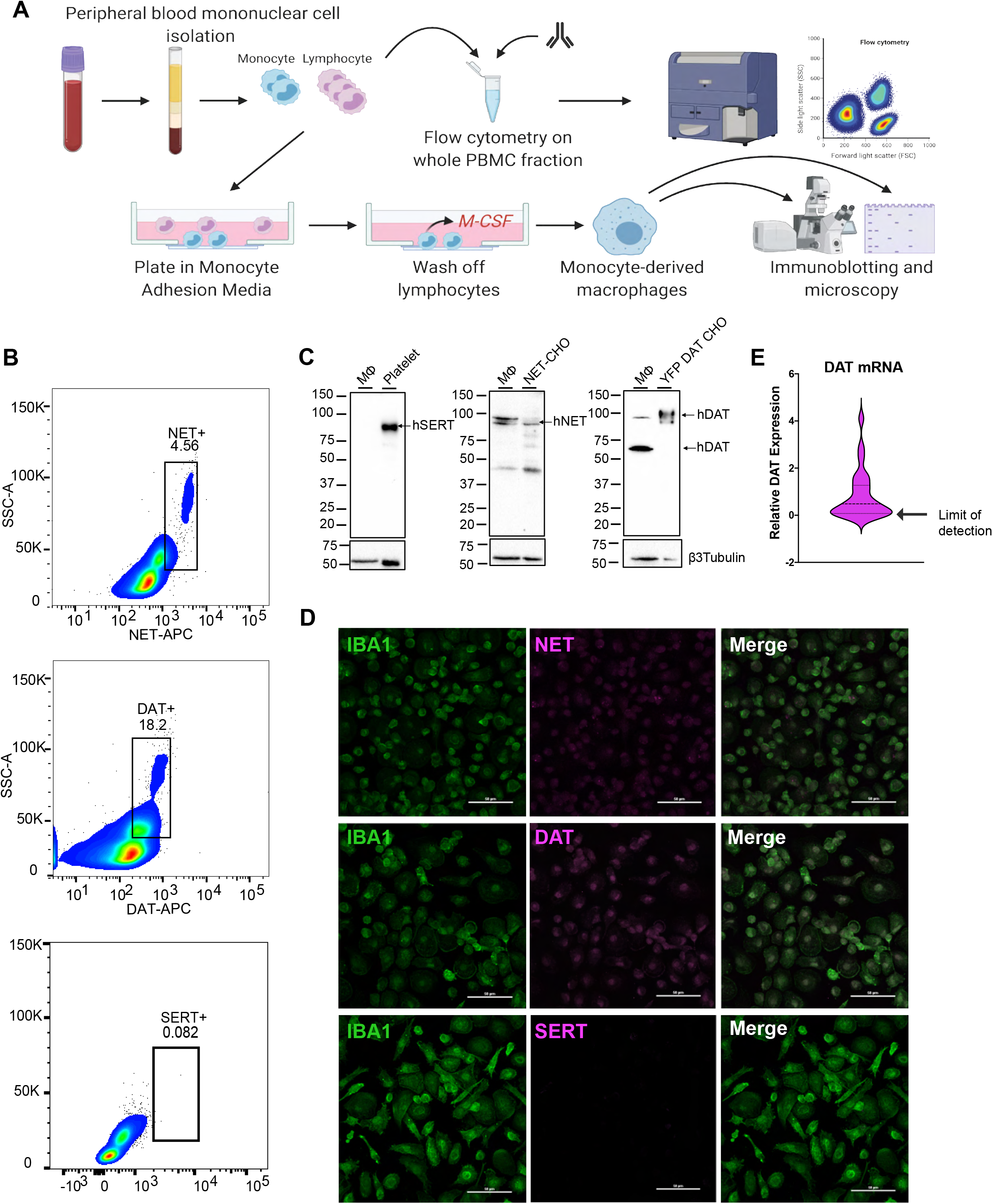
Human monocytes and monocyte-derived macrophages express NET and DAT, but not SERT. (A) Schematic depicting the isolation of PBMCs from human whole blood via density-dependent centrifugation with Ficol. A fraction of the isolated PBMCs were used for flow cytometric analysis and the remaining cells were plated in monocyte-adhesion media with autologous serum and allowed to differentiate into monocyte-derived macrophages. (B) Density plots of flow cytometry data on acutely isolated human PBMCs show that approximately 18.2% of monocytes are DAT+ and approximately 4.56% of monocytes are NET+ (scatter plots representative of 3 independent experiments). SERT was not detected on monocytes. (C) Representative western blots from lysates of cultured human monocyte-derived macrophages probed for SERT, NET, or DAT. Human monocyte-derived macrophages did not express SERT (positive control: human platelets) but did express both NET (positive control: NETexpressing CHO cells) and DAT (positive control: YFP-DAT-expressing CHO cells) (n=3 independent experiments). (D) Representative confocal images of human monocyte-derived macrophages immuno-stained for IBA1 and either NET, DAT, or SERT. IBA+ cells (macrophages) were positive for NET and DAT, but not SERT (n=3 independent experiments). E) RT-qPCR on cultured human monocyte-derived macrophages indicates that the mRNA for DAT protein is expressed in these cells (n=26).

As circulating monocytes are fundamentally different from engrafted macrophages in tissues, we next investigated whether MDMs retained NET and DAT expression after differentiation *in vitro*. To this end, we cultured monocytes into macrophages *in vitro* over 6-7 days (**Figure 1A**). Western blot analysis of MDM lysates confirmed that differentiated macrophages expressed NET (positive control, NET-expressing CHO cells), DAT (positive control, YFP-DAT-expressing CHO cells), but not SERT (positive control, human platelets, **Figure 1C**). These data are consistent with a recent study that showed expression of NET on murine and human adipose-tissue macrophages associated with sympathetic nerve terminals (Pirzgalska et al. 2017), however this study did not probe for DAT. Hence, the finding of DAT expression on human monocyte-derived macrophages is relatively novel. Therefore, we confirmed these findings using immunofluorescent staining and confocal microscopy. Cultured macrophages were labelled with Iba1, a pan-macrophage marker, and for NET, DAT, or SERT. Consistent with our western blot findings, Iba1+ cells showed signal for both NET and DAT but not for SERT (**Figure 1D**). Furthermore, qRT-PCR of cultured human macrophages revealed that the mRNA for DAT was also expressed in these cells (**Figure 1E**). Taken together, these data consistently support the interpretation that, human MDM express both NET and DAT, but not SERT.

### NET and DAT on human monocyte-derived macrophages is membrane-bound and functional

We next sought to determine if DAT or NET on human macrophages are functional. In order to do this, we first investigated the sub-cellular distribution of NET and DAT on cultured monocyte-derived macrophages, which are used to study live macrophage functions. Using cell surface biotinylation assay followed by immunoblotting, both NET and DAT were detected in the biotinylated membrane fraction of MDM at the appropriate molecular weights (**Figure 2A**). Notably, macrophages harbored an intracellular pool of DAT but not NET. To complement these data, we performed total internal reflective fluorescence microscopy (TIRF-M) which allowed us to visualize proteins at or near the basal membrane. We first validated our ability to detect surface DAT in TIRF-M using CFP-tagged DAT-expressing CHO cells, which showed robust overlap between the CFP tag and DAT immuno-labeling in the TIRF plane (**Supplemental Figure 2A**). Cultured macrophages were co-labelled with AlexaFluor-555 conjugated CTxB (CTxB-555), which binds the plasma membrane protein GM1, and with antibodies against NET or DAT. As expected, TIRF-M readily identified the plasma membrane marked CTxB-555 labeling. In concordance with our biotinylation data, scattered NET and DAT punctae were also detected in the TIRF plane, some of which co-localized with CTxB-555 punctae (**Figure 2B**). To confirm the TIRF-M data we employed Stimulated Emission Depletion (STED) super-resolution microscopy and confocal microscopy on macrophages co-labelled with CTxB-555 and anti-DAT antibody (with an AlexaFluor-647 secondary antibody). Both confocal (**Figure 2C**) and STED (**Figure 2D**) microscopies indicated modest co-localization between CTxB-555 and DAT, consistent with the abundant expression of GM1 at the plasma membrane and DAT localization to both the plasma membrane and intracellular compartment.

**Figure 2:**
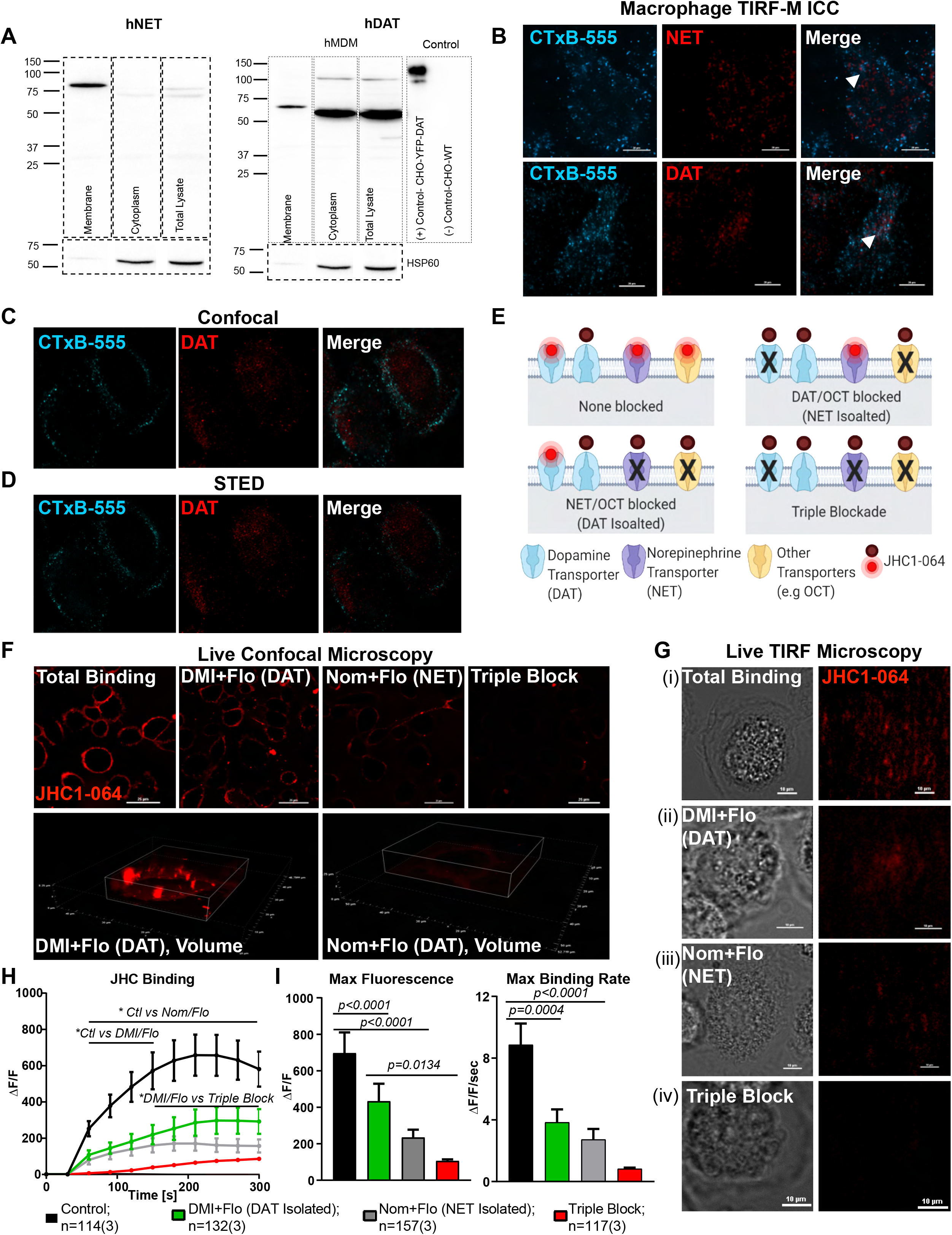
Human monocyte-derived macrophages express DAT and NET that are localized to the plasma membrane. A) Representative immunoblots for NET and DAT from macrophage intracellular and membrane fractions separated via biotinylation assay. The majority of NET in macrophages was localized to the membrane fraction, whereas DAT in macrophages localized to both the membrane fraction and the intracellular fraction (positive control: YFP-DAT expressing CHO cells). B) Representative images from Total Internal Reflective Fluorescence Microscopy (TIRF-M), which only visualizes the basal membrane, of cultured macrophages that were PFA-fixed, labelled with Alexafluor555-conjugated CTxB (a marker of membrane raft protein GM1) and either NET or DAT antibodies. CTxB555 punctae appeared on the basal membrane along with scattered punctae of NET (top) and DAT (bottom), indicating membrane localization. Some punctae of CTxB colocalized with NET or DAT (arrowheads, n=4 experiments), showing DAT and NET are distributed in both CTxB positive and CTxB negative loci at the membrane. (C and D) Representative confocal images (C, n=4 experiments) or Stimulated Emission Depletion (STED) super-resolution microscopy (D, n=2 experiments) on cultured human macrophages labelled with CTxB-555 and for DAT. Both modes of microscopy showed some of the DAT signal co-localizes with the CTxB-555 signal at or near the membrane. E) Cartoon schematic depicting the JHC1-064 binding assay by which JHC1-064, an analogue of cocaine, only fluoresces upon binding to outward-facing transporters such as DAT, NET, or related transporters. Using desipramine (DMI, NET antagonist), nomifensine (Nom, DAT antagonist), and fluoxetine (Flo, SERT and OCT antagonist) in different combinations allowed for selective isolation of transporter specific binding. (F) Live-cell confocal images of JHC1-064 binding to macrophages in the presence of no antagonists (total binding), NET/OCT blockade (DAT isolated binding), DAT/OCT blockade (NET isolated binding, or triple blockade (non-specific binding). (G) Live TIRF-M images of macrophages following JHC1-064 binding in the presence of absence of (i) no antagonists (total binding), (ii) NET/OCT blockade (DAT-isolated), (iii) DAT/OCT blockade (NET-isolated), (iv) or triple blockade shows progressive attenuation of JHC1-064 signal with the addition of the antagonists. (H) Quantifying the JHC1-064 signal from binding time-lapse images shown in (F) shows that blocking NET/OCT or DAT/OCT decreased JHC1-064 binding, and that blocking all 3 transporters further decreased the JHC1-064 signal on macrophages. (I) Blocking NET/OCT or DAT OCT decreased both magnitude (left) and rate (right) of JHC1-064 binding to human macrophages compared to control (Max. Fluorescence: Control vs DAT-isolated p<0.0001, control vs NET-isolated p<0.0001; Max Binding Rate: Control vs DAT-isolated p=0.0004, control vs NET-isolated p<0.0001). Blocking all three transporters further decreased the magnitude of JHC1-064 binding compared to the DAT-isolated condition (p=0.0134). Images and data in (F-I) are from n=114-157 cells/group from 3 independent experiments. Statistics in (I) were performed using a one-way ANOVA with Tukey’s post-hoc test.

To further validate these findings, we employed the use of JHC1-064, a membrane-impermeable fluorescent analogue of cocaine which only fluoresces upon binding to outward-facing biogenic amine transporters at the membrane (Cha et al. 2005). First, we validated our ability to measure JHC1-064-DAT binding in CHO cells transfected with CFP-DAT as a positive control. CFP is excited by the 405 nm laser line and detected at 485 nm, whereas JHC1-064 excitation and emission are 561nm and 617nm, respectively; therefore, there is a minimal bleed-through between the two channels. Live cell confocal and TIRF-M experiments in CFP-DAT expressing cells, showed overlay between the CFP–DAT signal and the JHC1-064 signal (**Supplemental Figure 2B,C**), demonstrating that JHC1-064 binds to DAT in these cells. The JHC1-064/DAT binding was blocked when the cells were pretreated with a DAT antagonist, nomifensine (10 µM), which we used as a negative control. The reason we did not use GBR12909 to block DAT is that unlike nomifensine, GBR12909 blocks both DAT and NET (**Table 1**). Therefore, for our studies we used nomifensine to identify DAT-specific activities.

**Table 1.**

| Table 1- Antagonist specificity/cross reaction | OCT/<br>Sigma1R | SERT | NET | DAT | References |
| --- | --- | --- | --- | --- | --- |
| **GBR ( DAT & NET blocker)** | Blocks | No effects | Blocks | Block | Heikkila RE and Manzino L. 1984; Husbands SH et al. 1999; Cao J. et al 2001 |
| **Fluoxetine (FLO)**<br>**(SERT/OCT blocker)** | Blocks | Blocks | No effect | No effect | Owens MJ et al. 1997 |
| **Desipramine (DMI)** | No effect | No effect | Blocks | No effect | Owens MJ et al. 1997 |
| **Nomifensine (NOM)** | No effect | No effect | No effect | Blocks | Rothman R.B. et al. 1993 |

The DAT/JHC1-064 binding data in the CHO-DAT cells with and without nomifensine were used as a positive and negative control groups to measure JHC1-064 binding to human macrophages. *Live-cell microscopy required us to study NET and DAT activity separately. FACS sorting of live monocytes that were either NET+ or DAT+ (i.e., not double-positive), was not possible since these antibodies bind intracellular epitopes and would require fixation. Additionally, it is possible that DAT or NET expression change over the culturing period. Therefore, to isolate NET and DAT activity, we studied DAT or NET function in the presence of NET antagonists to isolate DAT-specific activity and vice versa (**Figure 2E**).* Via live-cell confocal (**Figure 2F**) and TIRF microscopy (**Figure 2G**) we measured JHC1-064 binding in human macrophages in 4 different conditions: 1) no antagonists present, control condition representing total binding; 2) pretreatment with fluoxetine (Flo, 5 µM) and nomifensine (Nom, 10 µM) to selectively block DAT and OCTs which isolates NET/JHC1-064 binding; 3) pretreatment with fluoxetine and desipramine (DMI, 10µM) to block OCTs, and NET, which isolates DAT/JHC1-064 binding; and 4) pretreatment with fluoxetine, desipramine, and nomifensine which blocks all specific binding. As predicted, the highest JHC1-064 binding was detected when no antagonists were present. The magnitude of JHC1-064 binding (**Figure 2H, I, left,** n=114-157 cells from 3 experiments/group) was attenuated when macrophages were pretreated with fluoxetine + desipramine (p<0.0001) or fluoxetine + nomifensine (p<0.0001), and further attenuated when all three antagonists were present (p=0.0134). A similar pattern was observed in the rate of JHC1-064 binding (**Figure 2I, right**), although not all the differences reached significance. These same trends for NET/JHC1-064 binding and DAT/JHC1-064 binding were qualitatively observed in TIRF-M (**Figure 2G**). While the detection of NET activity on human macrophages was anticipated, identifying the nomifensine sensitive DAT activity on these cells was unexpected. Overall, these data support the conclusion that both DAT and NET are expressed on the plasma membrane of human macrophages, they can bind to JHC1-064 can specifically identify the activity of each transporter individually using this pharmacological approach.

Both NET and DAT canonically work to uptake extracellular substrate into the intracellular space. Therefore, we next asked if NET and DAT on cultured human macrophages were capable of uptake. To this end, we used IDT307, a membrane-impermeable substrate for neurotransmitter transporters that only fluoresces after entering the cell (Blakely et al. 2011), to assay macrophages for transporter uptake (**Figure 3A**). First, we confirmed our ability to measure DAT specific uptake by quantifying IDT307 fluorescence in CFP-DAT expressing CHO cells in the presence of vehicle (positive control) or nomifensine (10µM, negative control, **Supplemental Figure 3A**). As expected, CFP-DAT CHO cells readily took up the IDT307 and fluoresced, and nomifensine reliably abolished uptake and fluorescence (**Supplemental Figure 3B, C**, p<0.0001, n=19-27 cells/group from 3 experiments). We repeated this assay in macrophages treated with the same 4 conditions described above—no antagonists, pharmacological isolation of NET activity (Nom + Flo), pharmacological isolation of DAT activity (DMI + Flo), and triple antagonist treatment (**Figure 3B,** n=88-294 cells/group from 3 experiments). Similar to the JHC1-064 assay, we observed the most IDT307 uptake without any antagonists. The magnitude and rate of IDT307 uptake was attenuated by both the nomifensine + fluoxetine (Nom + Flo, AUC p<0.0001, Average slope p<0.0001) and desipramine + fluoxetine (AUC p<0.0001, Average slope p<0.0001) conditions. Addition of all three antagonists—simultaneously blocking DAT, NET and other transporters—further attenuated uptake (**Figure 3C**, AUC DAT vs Triple p=0.0008, AUC NET vs Triple p=0.0016, Average slope DAT vs Triple p<0.0001, Average slope NET vs Triple p<0.0001). Subtracting fluorescence of the triple antagonist condition from either the NET-isolated or DAT-isolated conditions revealed notable and separate NET-specific (**Figure 3D**) and DAT-specific (**Figure 3E**) uptake capacities of human macrophages, respectively. These data indicate that both NET and DAT on human monocyte-derived macrophages can uptake substrate from the extracellular milieu.

**Figure 3:**
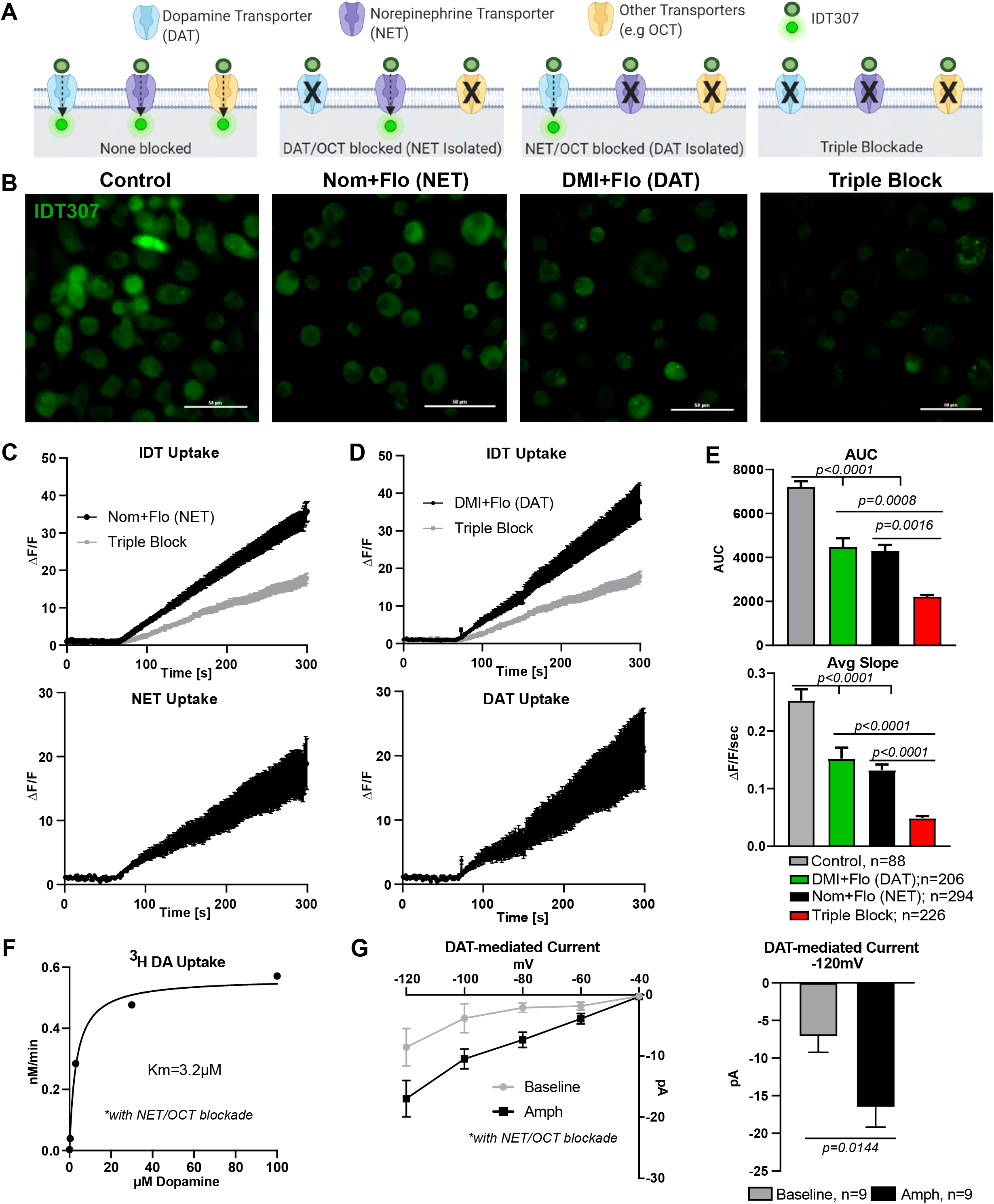
NET and DAT on human macrophages can work in uptake-mode. (A) Schematic cartoon depicting the experimental design employing various conditions to measure the uptake of IDT307, a synthetic substrate for neurotransmitter transporters which fluoresces upon entry into the cell. Uptake of IDT307 was assayed under four conditions to determine transporter specific uptake of IDT307: 1) with no antagonists (showing total uptake); 2) with DAT/OCT blocked (putatively isolating NET-mediated uptake); 3) with NET/OCT blocked (putatively isolating DAT-mediated uptake); and 4) triple blockade of DAT/NET/OCT to eliminate transporter-specific uptake. (B) Representative images following perfusion with IDT307 show that human macrophages exhibited bright fluorescence. Fluorescence was decreased with addition of either DAT/OCT blockers and NET/OCT blockers, and fluorescence was further attenuated in the presence of triple blockade. (C) IDT307 uptake measured as cellular fluorescence above baseline in the triple blockade condition was subtracted from the NET-isolated condition (top) to give the NET-mediated IDT307 uptake (bottom). (D) IDT307 uptake measured as cellular fluorescence above baseline in the triple blockade condition was subtracted from the DAT-isolated condition (top) to yield the DAT-mediated IDT307 uptake (bottom). (E) Quantifying the magnitude (AUC, top) and average slope (bottom) of IDT307 uptake by human macrophages across the four conditions confirms that blockade of either NET/OCT or DAT/OCT significantly decreases the magnitude and rate of IDT uptake (AUC: Control vs DAT-isolated p<0.0001, control vs NET-isolated p<0.0001; Average slope: Control vs DAT-isolated p<0.0001, control vs NET-isolated p<0.0001), and the triple antagonist cocktail further decreases the magnitude and rate of IDT uptake (AUC: Control vs DAT-isolated p=0.0008, control vs NET-isolated p=0.0016; Average slope: Control vs DAT-isolated p<0.0001, control vs NET-isolated p<0.0001). Images and data in (B-E) are from n=88-294 cells/group from 3 independent experiments, and statistics were performed using a one-way ANOVA with Tukey’s post-hoc test. (F) Nomifensine-sensitive uptake of tritiated dopamine (^3^H-DA) by cultured human macrophages in the presence of a NET/OCT blockade shows a first-order uptake kinetics shown as nM/min with a Km of approximately 3.2μM. Data from 2 independent experiments. (G) DAT-mediated inward currents on cultured human macrophages were recorded via whole-cell patch clamp in the voltage-clamp mode. Serial hyperpolarizing steps evoked families of inward currents which were augmented by DAT activator amphetamine (Amph) and blocked by nomifensine (Nom) (left). Amphetamine induced significantly increased DAT-mediated inward current at -120mV (right, p=0.0144, unpaired t-test). Data are from 9 experiments/group.

While NET-mediated uptake on adipose-resident macrophages has previously been shown (Pirzgalska et al. 2017), the finding of a separate DAT-mediated uptake was surprising. Therefore, we employed two additional complementary approaches to validate our findings. First, we used the classical tritiated dopamine uptake assay and found that human MDM exhibited Michaelis-Menten-like kinetics for dopamine uptake, with a K_m_ of approximately 3.2 μM (**Figure 3F**). This is similar to the K_m_ observed in other DAT-expressing systems (Mazei-Robison and Blakely 2005). In addition, DAT-mediated substrate uptake is coupled to the co-transport of 2 Na^+^ ions and one Cl^−^ ion, generating a net-inward current that can be measured via whole cell voltage-clamp electrophysiology by serial hyperpolarizing voltage steps. Mirroring our positive control cells (stably transfected YFP-DAT expressing HEK cells, **Supplemental Figure 3D**), human MDM showed a nomifensine-sensitive inward current, albeit with a lower magnitude, that was increased following DAT activation with amphetamine (**Figure 3G,** p=0.0144). Altogether, our findings indicate that both NET and DAT are expressed at the membrane of primary human monocyte-derived macrophages and function in their canonical uptake mode in these cells. Next, we investigated whether DAT expression in limited to circulating monocytes and MDM or other tissue-resident macrophages also express DAT.

### A portion of human gut-resident macrophages are DAT+

A recent study reported that human sympathetic-nerve associated adipose macrophages express a functional NET (Pirzgalska et al. 2017); however, to our knowledge, DAT expression has not been thoroughly investigated on any tissue-resident macrophage population. While our data indicate that human monocytes and monocyte-derived macrophages *in vitro* express DAT, cultured macrophages lack the tissue-derived signaling factors and thus do not faithfully recapitulate the phenotype of any tissue-resident macrophage (Gosselin et al. 2014). Thus, before proceeding, we asked if any human tissue-macrophage populations expressed DAT *in situ*. The gut is a rich source of dopamine (Matt and Gaskill 2019), and its resident macrophage pool is partially maintained by monocyte-derived macrophages (Bain et al. 2014). Therefore, we hypothesized that human gut-resident macrophages express DAT.

We curated a large single-cell RNA sequencing data set on biopsy samples of human colon from the NCBI Gene Expression Omnibus (Li et al. 2020). Clustering analysis of this data set using the Seurat pipeline as previously described (Satija et al. 2015) yielded 13 transcriptionally distinct clusters which we annotated as various epithelial, muscle, and immune populations based on their differentially expressed genetic markers (**Figure 4A**). Searches for expression of the DAT gene (SLC6A3) showed low expression levels in several clusters, notably cluster 7. Cluster 7 was also enriched in markers such as AIF-1, CD206, CD163, CYBB, CD86, and IL-10, consistent with the identity of tolerizing gut-resident macrophages (**Figure 4B**). Thus, these data suggest that, in addition to monocytes and cultured monocyte-derived macrophages, the DAT is expressed in at least some human gut-resident macrophages.

**Figure 4:**
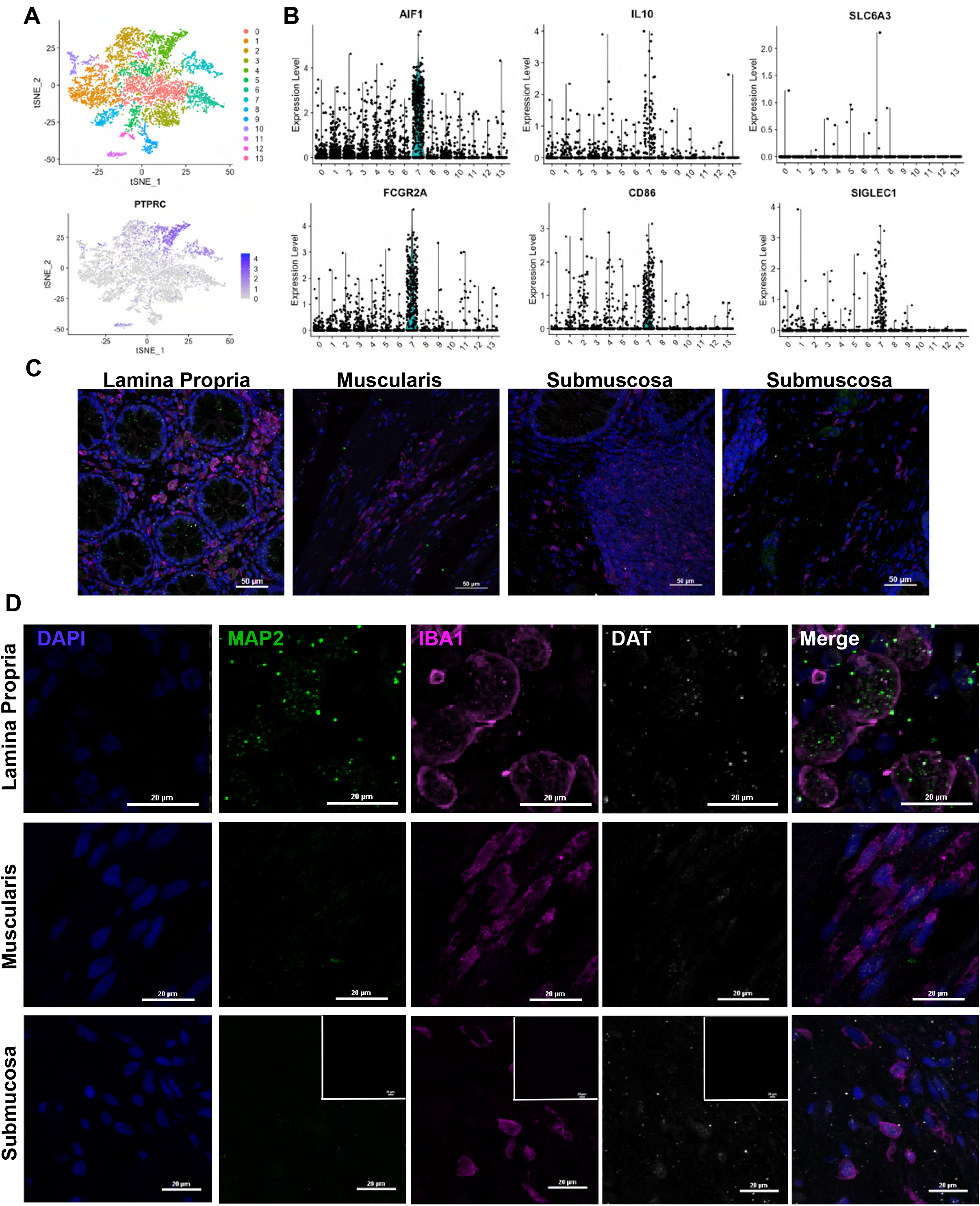
A subpopulation of human intestinal macrophages express DAT. (A) A previously published single-cell RNA sequencing data set was procured from NCBI’s Gene Expression Omnibus (GEO) using the search terms “gut” and “macrophage”. The data set was analyzed using the R package Seurat to cluster the cells based off the 10 most significant principal components and dimensionally reduce the data using t-stochastic-neighbor embedding (t-sne) plots yielding 13 different clusters of cells (top). Relative expression of PTPRC (CD45) was overlaid on the t-sne plot (bottom). (B) Relative expression of macrophage markers AIF1/IBA1, FGCGRA, IL-10, CD86, and SIGLEC1 in addition to expression of SLC6A3 (DAT) in each of the 13 clusters represented as violin plots indicate that cluster 7 was enriched for macrophage markers and contained some SLC6A3-expressing cells. (C) Representative confocal microscopy images of healthy human colon tissue labelled for IBA1 (macrophages), MAP2 (neurons), DAPI (nuclei), and DAT. Images were collected from various anatomical parts of the gut wall including the lamina propria (left), muscularis (middle-left), submucosa containing gut-associated lymphoid tissue (middle-right), and submucosa containing neuronal ganglia (right). All areas contained IBA1+ cells (macrophages). (D) High magnification images from each of the anatomical regions shown in (C). Lamina propria contained isolated IBA1+ cells and IBA1+ cells enveloping MAP2+ areas. Some, but not all submucosal IBA1+ cells were weakly DAT+, whereas muscularis macrophages were DAT-. Secondary only negative controls shown as inset in the bottom of (D). Images in (C&D) are from 3 independent experiments from 1 healthy human donor.

An important limitation of the single-cell transcriptomic approach of a diverse sample population is the low sensitivity to low-abundance transcripts. To validate DAT protein expression in human gut-resident macrophages, we complemented the scRNA-seq analysis with confocal microscopy in situ. In the gut, macrophage heterogeneity is partially governed by their anatomical niche within the intestinal wall (De Schepper et al. 2018, Asano et al. 2015, Gabanyi et al. 2016, Hadis et al. 2011), therefore, we examined macrophage populations in the human colon lamina propria, submucosa, and muscularis. Tissues were immunolabeled for Iba1 (pan-macrophage marker), MAP2 (neuronal marker) and DAT, and assessed using confocal microscopy (**Figure 4C**). Iba1+ cells were abundant in all locations. In the lamina propria, Iba1+ cells enwrapping MAP2+ puncta were DAT+, but it was difficult to determine whether the DAT signal was present on neurons or macrophages, or both (**Figure 4D, top**). However, in the submucosa, we found a subgroup of macrophages that were Iba1+/DAT+(**Figure 4D, bottom**, secondary only negative control inset). Interestingly, most of the Iba1+/DAT+ cells were found near lymphoid-like follicles or MAP2+ ganglia, but did not colocalize with MAP2+ areas, suggesting this niche may be associated with DAT expression in gut macrophages. These data indicate that in addition to circulating monocytes and cultured human monocyte-derived macrophages, human gut-resident macrophages also express DAT *in situ*. Next, we investigated whether or not DAT activity modulates macrophage immune functions such as cytokine secretion and phagocytosis.

### DAT activity modulates macrophage immune functions

Since our data indicate that human MDM macrophages express a functional DAT (**Figures 1-4**), we next asked if DAT activity affected on macrophage immune functions such as cytokine secretion and phagocytosis. We first investigated whether or not DAT activity modulates monocyte and macrophage cytokine profile. To this end, we used lipopolysaccharide (LPS), a product of gram-negative bacterial cell walls that agonizes CD14/TLR4 and elicits a potent inflammatory response. Since DAT activators, such as amphetamine and DA have both DAT and non-DAT dependent effects on immune function, we first chose to simply inhibit DAT activity with nomifensine. Therefore, we treated monocytes (n=6) and monocyte-derived macrophages (n=11-12/group) with either vehicle, nomifensine, LPS, or LPS + nomifensine, all in the presence of NET blockade for 24 hours. In cultured macrophages LPS significantly increased release of pro-inflammatory cytokines IL6 (p=0.0055), TNFα (p=0.0041), and CCL2 (p=0.0019) in MDM. The effects of LPS on IL6 and TNFα were significantly increased in the presence of a DAT blockade (**Figure 5A**, IL6 p=0.0216, TNFα p=0.0030). In the absence of LPS, DAT blockade had no effect on the baseline release of these cytokines. We observed a similar effect in freshly isolated monocytes, with DAT blockade enhancing LPS-induced production of intracellular cytokines (**Supplemental Figure 4**). Notably, DAT inhibition had no effect on LPS-induced release of IL1β (**Supplemental Figure 5A**), and NET inhibition had no significant effect on LPS-induced release of IL6, TNFα, or CCL2 (**Supplemental Figure 5B**). Thus, DAT activity can modulate the cytokine response independent of NET activity.

**Figure 5:**
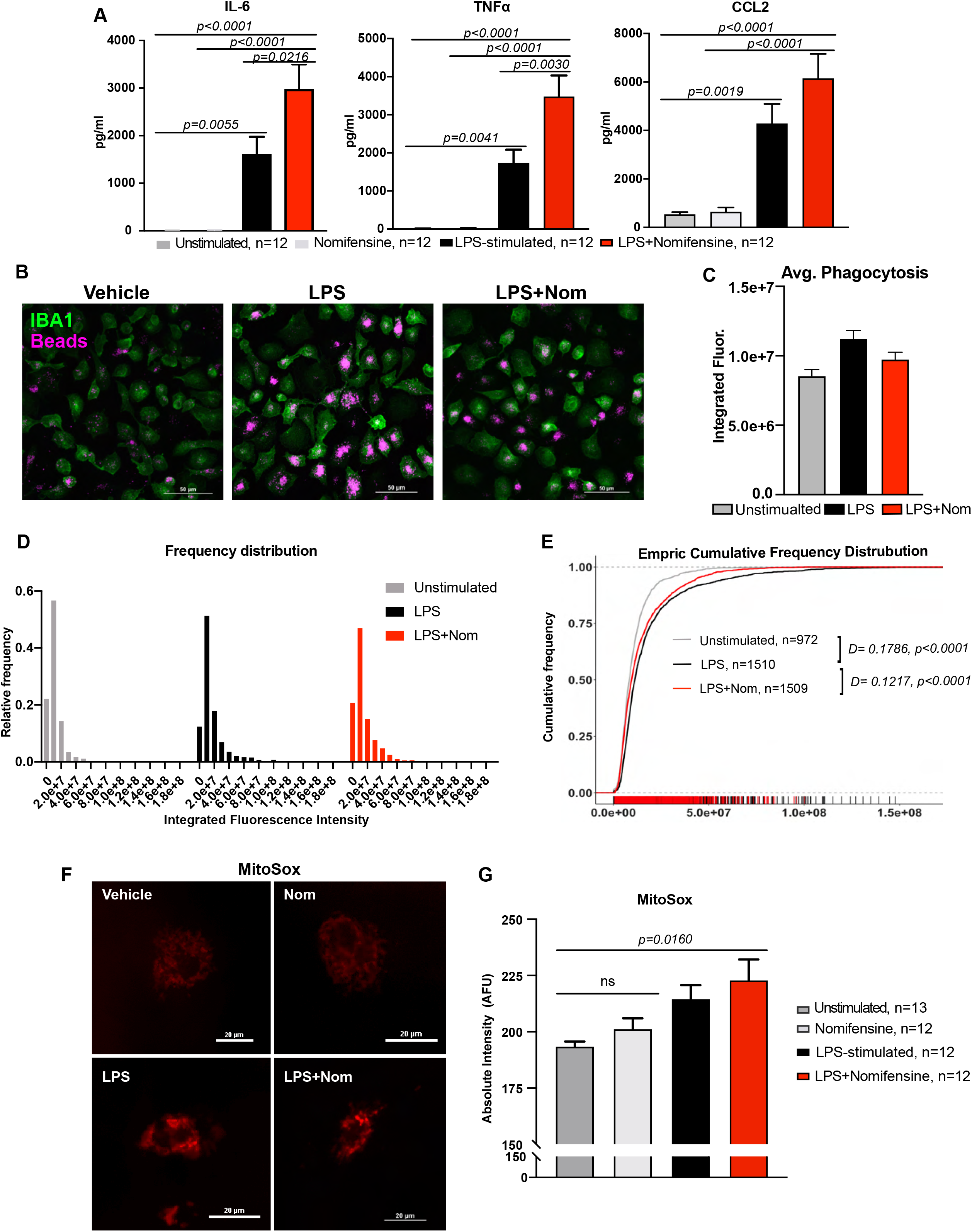
Inhibition of DAT enhances the pro-inflammatory program in response to LPS. (A) Cultured human macrophages were treated with vehicle, nomifensine, LPS, or LPS+ nomifensine in the presence of NET/OCT blockade and their conditioned media was collected for measurement of IL-6, TNF-α, and CCL2. LPS treatment significantly increased the secretion of all 3 soluble factors (IL-6 p=0.0055, TNFα p=0.0041, CCL2 p=0.0019). While nomifensine alone had no effect on cytokine/chemokine secretion, nomifensine in the presence of LPS significantly increased the LPS-induced secretion of TNFα (p=0.0030) and IL6 (p=0.0216) and had a similar effect on CCL2 secretion. Data are from n=12 experiments/group, and statistics were performed using a one-way ANOVA with Tukey’s post-hoc test. (B) Representative images of cultured human macrophages treated with vehicle (media), LPS, or LPS+ nomifensine and incubated with fluorescent latex beads, fixed and labelled for IBA1, and then imaged using confocal microscopy to measure phagocytosis. (C) The average integrated fluorescence intensity of phagocytic beads within macrophages, indicative of average phagocytic capacity, showed a slight increase in LPS and a non-significant decrease with LPS+ nomifensine. (D) Frequency histograms showing the skewed distribution of fluorescence intensity of phagocytic beads in macrophages across the 3 conditions. (E) Empiric cumulative frequency distribution curves, of unstimulated, LPS-stimulated, and LPS+ nomifensine-treated macrophages shows that LPS significantly increased phagocytosis compared to unstimulated condition (D-statistic=0.1786, p<0.0001), and co-treatment with LPS and nomifensine decreased macrophage phagocytosis back towards unstimulated levels (D-statistic=0.1217, p<0.0001 vs LPS-stimulated). Images and data in (B-E) are from n=972-1510 cells/group across 3 independent experiments. Statistics were performed using K-S tests. (F) Representative images of cultured human macrophages treated as in (A) and incubated with a red-shifted fluorescent indicator of mitochondrial superoxide species (MitoSox Red™). (G) While treatment with LPS slightly, but not significantly, increased mitochondrial superoxide levels, as measured by increased fluorescence intensity, only co-treatment with LPS and nomifensine produced a significant increase in mitochondrial superoxide levels compared to vehicle (p=0.0160). Images and data in (F&G) are from n=12-13 experiments/group, and statistics were performed using a one-way ANOVA with Tukey’s post-hoc test.

We next asked if DAT blockade had a similar effect on macrophage phagocytosis. Blocking NET, we utilized the experimental approach described by the Tsirka lab to quantify phagocytosis of fluorescent latex beads by Iba1+ macrophages (**Figure 5B, C**) via a publicly available pipeline for analysis (Caponegro et al. 2020). The analysis we employed relies on optimal mass transport theory and utilizes empiric cumulative distribution frequencies rather than measures of central tendency to detect shifts in phagocytosis capacity of macrophage populations. When applied to our conditions, we detected a rightward shift in phagocytosis induced by LPS (D-statistic = 0.1786, p<0.0001), consistent with increase in phagocytosis capacity. Blockade of DAT in the presence of LPS resulted in a leftward shift, bringing phagocytic capacity back towards baseline levels (**Figure 5D, E**, D-statistic =0.1217, p<0.0001). Importantly, these data only assess two macrophage functions and do not provide a comprehensive picture of the macrophage phenotype. Nevertheless, the increased pro-inflammatory cytokine release and decreased phagocytosis induced by DAT blockade suggest loss of DAT activity skews macrophages towards a more pro-inflammatory state.

Mitochondrial health and mitochondrial oxidative stress are closely associated with macrophage phenotype and inflammatory function (Mills, Kelly and O’Neill 2017, Minhas et al. 2019). Therefore, we next investigated whether or not DAT blockade during LPS-stimulation affected mitochondrial oxidative stress. Macrophages were treated with MitoSox Red™, a fluorescent reporter of mitochondrial superoxide species (**Figure 5F,** all conditions contained NET blockade) during LPS and nomifensine treatment. Surprisingly, while LPS stimulation increased mitochondrial oxidative stress, this increase was not significant and only DAT blockade in the presence of LPS produced a significant elevation in mitochondrial superoxide levels (**Figure 5G**, p=0.0160). Although these data do not directly assess mitochondrial function, they support the notion that DAT blockade may affect mitochondrial health during inflammation via oxidative burden. Taken together, these data implicate DAT activity as an important immunomodulator during the immune response to LPS.

### LPS decreases DAT-mediated uptake without changing DAT membrane levels

To further examine the interaction between DAT activity and the macrophage inflammatory response, we examined the direct effects of LPS on DAT function, with the hypothesis that LPS directly alters DAT activity. Literature on characterizing DAT activity and regulation in neurons and heterologous cells indicates that DAT activity is regulated by several mechanisms including trafficking to and from the plasma membrane and changes in its conformational state (Vaughan and Foster 2013). One well-characterized regulator of DAT activity is PKC-induced internalization of DAT and subsequent decrease in DAT-mediated uptake (Melikian and Buckley 1999). Before investigating potentially novel LPS-induced regulatory mechanism of DAT activity, we first tested whether or not DAT on macrophages was subject to the well-characterized PKC-induced internalization. To this end, via live-cell TIRF-M, we monitored membrane DAT-JHC1064 complexes in both macrophages and YFP-DAT expressing cells before and after addition of PKC-activator phorbol myristate acetate (PMA, 1µM) in the presence of a NET blockade. Treatment with PMA decreased the TIRF-M footprint in both YFP-DAT cells (positive control group, p=0.0023) and macrophages (p=0.0015), indicative of DAT internalization in both cell types (**Supplemental Figure 6A, B**). In parallel experiments, we examined whether PMA treatment could alter DAT-mediated IDT307 uptake in macrophages in the presence of NET blockade. Live-cell imaging revealed that activating PKC using PMA significantly decreased the DAT-mediated uptake of IDT307 human macrophages (**Supplemental Figure 6C-E**), consistent with DAT internalization. Together, these data indicate that DAT on macrophages undergoes canonical PKC-induced internalization, and that we can use microscopy to accurately detect changes in DAT surface expression and activity on macrophages.

We next shifted our focus to possible LPS-induced regulation of DAT activity. We repeated the IDT307 uptake assay in unstimulated macrophages and macrophages treated with LPS for 24 hours (**Figure 6A**, both conditions contained NET blockade). Surprisingly, we observed a dramatic decrease in DAT-mediated uptake of IDT307 in the LPS-stimulated macrophages (**Figure 6B**) as both the magnitude (AUC, p<0.0001) and rate (Average slope, p=0.0219) of IDT were decreased by LPS stimulation (**Figure 6C**). Our previous data indicate that DAT-mediated uptake is accompanied by a nomifensine-sensitive inward current in heterologous cells and on macrophages (**Supplemental Figure 3D, Figure 3G**). Therefore, to validate our findings from the IDT uptake experiment, we performed patch clamp recordings in whole-cell configuration on unstimulated and LPS-stimulated macrophages with NET blocked. In unstimulated macrophages, we reproduced our earlier findings and measured a nomifensine-sensitive, amphetamine-induced inward current that was significantly decreased by LPS stimulation (**Figure 6D, E**, p=0.0084, p=0.0038). Taken together, these findings support the hypothesis that LPS stimulation decreases DAT-mediated uptake.

**Figure 6:**
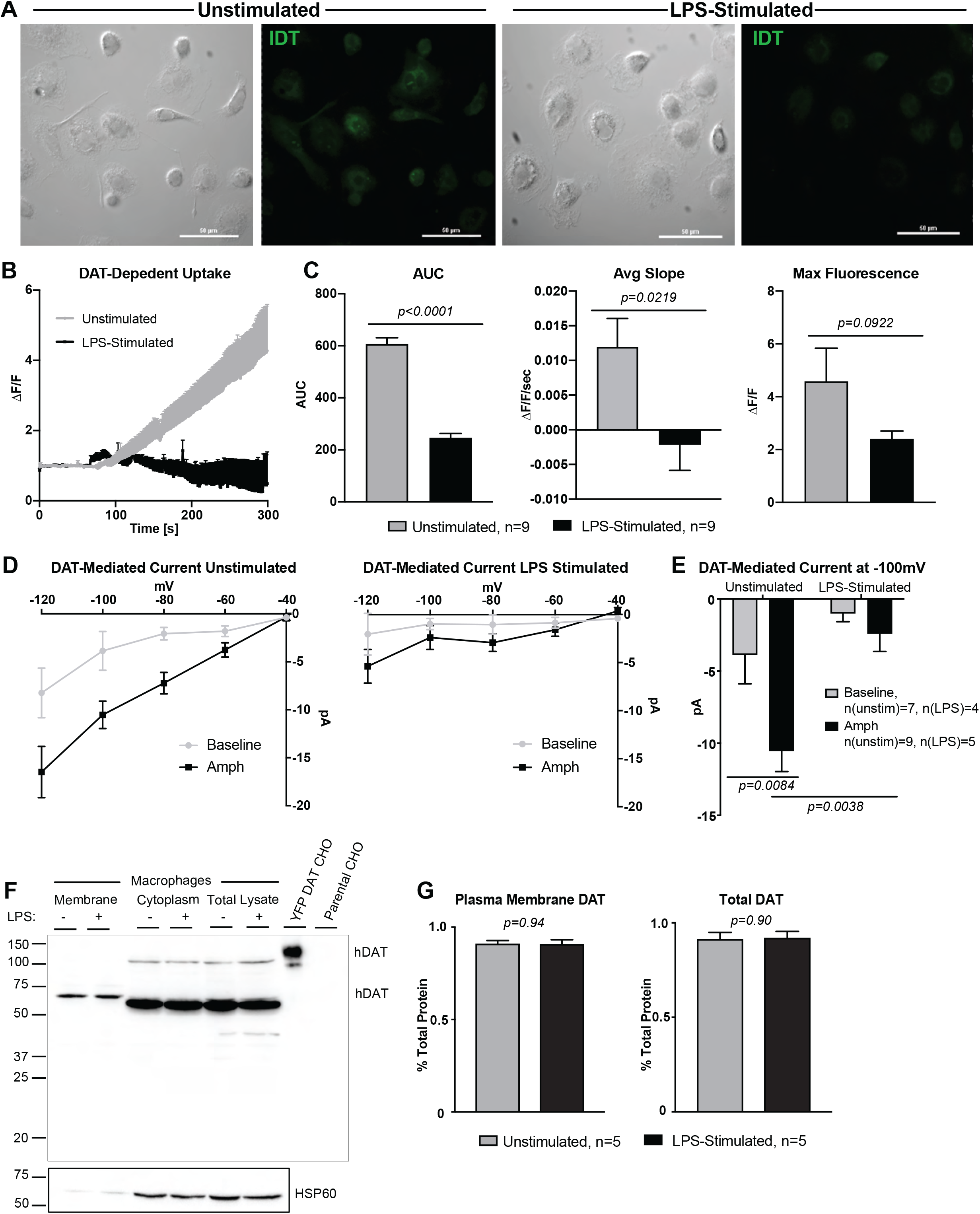
LPS stimulation decreases DAT-mediated uptake without affecting DAT membrane-levels. (A) Cultured human macrophages were unstimulated or LPS-stimulated and assayed for DAT-mediated uptake by IDT307 perfusion in the presence of NET/OCT block under live-cell imaging as in Figure 3. (B) DAT-mediated uptake for both LPS-stimulated and unstimulated macrophages was calculated by blocker-subtractions as in Figure 3 and showed that LPS-stimulated macrophages exhibited decrease DAT-mediated uptake of IDT307 compared to unstimulated controls. (C) Quantifying the uptake in (B) shows that LPS-stimulation decreased both the magnitude (AUC, p<0.0001 unpaired t-test; Max Fluorescence, p=0.0922 unpaired t-test) and the rate (Average slope, p=0.0219 unpaired t-test) of DAT-dependent IDT307 uptake. Data and images are from n=9 independent experiments/group. (D) Cultured human macrophages were either unstimulated (left) or LPS-stimulated (right) and patch clamped in whole-cell configuration under voltage clamp mode in the presence of NET/OCT block. DAT-mediated inward currents were calculated by subtracting the inward current in the presence of nomifensine from the inward currents either at baseline (baseline) or in the presence of DAT-activator, amphetamine (Amph). (E) DAT activation with amphetamine induced a significantly increased nomifensine-sensitive inward current (p=0.0084, two-way ANOVA with Sidak’s multiple comparison test); however, LPS-stimulated significantly decreased the amphetamine-induced DAT-mediated current (p=0.0038, two-way ANOVA with Sidak’s multiple comparison test). Data in (D-E) are from 4-9 experiments/group (specified in figure). (F) Representative western blot for DAT on membrane, cytosolic, and whole lysate fractions of unstimulated and LPS-stimulated human macrophages separated via biotinylation with YFP-DAT overexpressing CHO cells as positive control and parental CHO cells as negative control. The presence of HSP60 in the cytosolic and total, but not membrane fractions indicates successful fractionation. DAT bands were noted at the appropriate molecular weight (∼63kDa and ∼110kDa) in all lanes except negative control. (G) Densitometric quantification revealed LPS had no effect on membrane-localized DAT (left, p=0.94, unpaired t-test) or total DAT (right, p=0.90, unpaired t-test). Blot and data in (F-G) are from n=5 independent experiments/group.

Decreases in DAT uptake and inward current can reflect decreases in membrane-localized DAT, decreases in total DAT, changes in outwardly vs. inwardly conformational state of DAT, altered transport kinetics, or a combination thereof. Therefore, we investigated whether the reduced DAT-mediated uptake induced by LPS was due to decreased DAT membrane localization. Unstimulated and LPS-stimulated macrophages were biotinylated and lysed to separate membrane and intracellular fractions, and then each fraction was probed for DAT by immunoblot. Lysates from YFP-DAT expressing CHO cells were used as a positive control and lysates from parental CHO cells were the negative control (**Figure 6F**). DAT was detected in both membrane and cytosolic fractions in both conditions, and no difference was observed in either membrane DAT (p=0.94) or total DAT (p=0.90) between conditions (**Figure 6G**). Collectively, these findings indicate that LPS induces a decrease in DAT-mediated uptake without affecting total or surface DAT levels.

### LPS stimulation favors an efflux-promoting DAT

Our lab and others have previously shown that in addition to uptake, DAT can engage in reverse transport—termed efflux—a process by which it transports the substrate into the extracellular space (Khoshbouei et al. 2003, Khoshbouei et al. 2004, Sambo et al. 2017). DAT-mediated efflux is associated with drugs of abuse (Kahlig et al. 2005), ADHD (Mazei-Robison et al. 2008, Hansen et al. 2014), and autism (Bowton et al. 2014). Since LPS decreased DAT-mediated uptake without affecting membrane DAT levels, we hypothesized that LPS might alter DAT activity to favor efflux over uptake. We tested this using simultaneous patch-clamping and amperometry (Khoshbouei et al. 2003, Gnegy et al. 2004, Binda et al. 2008). In this technique, cells are patch clamped in whole-cell configuration in voltage-clamp mode, and the intracellular compartment is dialyzed with dopamine via the patch pipette. Simultaneously, an amperometry electrode is placed adjacent to the cell and held at 700 mV positive potential, which is above dopamine’s oxidation potential. In the presence of known efflux inducers, such as amphetamine, depolarizing steps promote a nomifensine-sensitive dopamine efflux. As the dopamine leaves the cell through the DAT, it binds to the amperometry electrode, causing it to oxidize and subsequently generate a detectable current (**Figure 7A**). We used this technique to measure DAT-dependent dopamine efflux in the presence of a NET blockade in both unstimulated and LPS-stimulated macrophages. We did not detect dopamine efflux at baseline in unstimulated macrophages; however, LPS-stimulated macrophages exhibited a significantly higher DAT-dependent dopamine efflux at baseline (**Figure 7B, top**, p=0.0243). Notably, LPS had no further effect on the amphetamine-induced efflux (**Figure 7B, bottom**, p=0.9356) overall indicating that LPS promotes DAT-mediated efflux on macrophages. To our knowledge, this is the first report of a physiologic case of DAT-dependent efflux, as this function has traditionally been studied in cases of genetic mutations or exogenous drugs.

**Figure 7:**
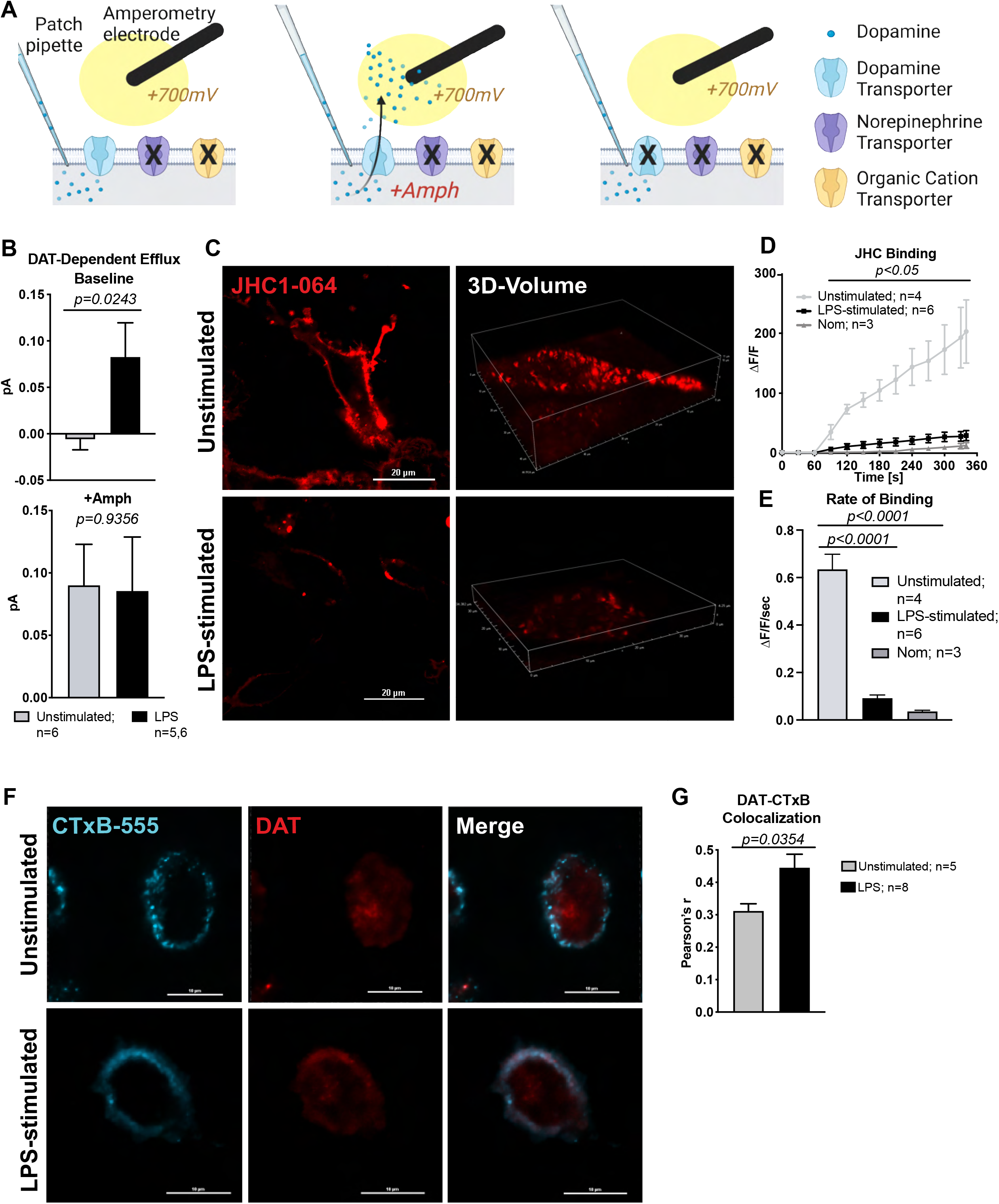
LPS-stimulation induces an efflux-promoting DAT on human macrophages. (A) Cartoon schematic representation of the simultaneous whole-cell patch-clamp and amperometry technique that measures dopamine efflux with msec resolution. In the presence of NET/OCT blockade, cells were patched in whole-cell configuration and dialyzed with dopamine via the patch pipette. Simultaneously, an amperometry recording electrode was held adjacent to the cell membrane at +700mV. The cell was depolarized via the patch electrode. In the presence of a known DAT efflux-inducers (e.g., Amphetamine), depolarization causes DAT-mediated efflux of dopamine, which is oxidized by the high electric potential generating a current measured via the amperometry electrode. In the presence of a DAT blocker, such as nomifensine, depolarization does not produce any efflux and no current is measured. DAT-mediated efflux was calculated by subtracting the current measured in the nomifensine condition from the currents measured at baseline or in the presence of amphetamine (Amph). (B) Unstimulated and LPS-stimulated human macrophages were subjected to the experimental setup in (A). While unstimulated macrophages did not exhibit any measurable DAT-mediated efflux at baseline, LPS-stimulation significantly increased basal DAT-mediated dopamine efflux (top, p=0.0243, unpaired t-test), but did not further affect the amphetamine-induced dopamine efflux (bottom, p=0.9356, unpaired t-test). Data are from n=4-6 experiments/group. (C) Representative images of JHC1-064 binding to unstimulated and LPS-stimulated macrophages in the presence of NET/OCT blockade taken using confocal microscopy, showing notably decreased JHC1-064 fluorescence at the plasma membrane in LPS-stimulated macrophages. (D) Quantifying fluorescence signal showed that LPS significantly decreased the magnitude of JHC1-064 binding to levels similar to those seen with DAT-blockade (nomifensine-treated, p<0.05). (E) The average rate of JHC1-064 binding (average slope from D) was similarly decreased by LPS-stimulation (p<0.0001, one-way ANOVA with Tukey’s post-hoc test) and by presence of DAT-specific antagonist, nomifensine (p<0.0001, one-way ANOVA with Tukey’s post-hoc test). Images and data in (D-E) are from n=3-6 independent experiments/group. (F) Representative confocal images of fixed unstimulated and LPS-stimulated macrophages labelled for GM-1 (a membrane marker) using CTxB-555 and DAT. CTxB-555 localized at or near the membrane whereas the DAT signal was distributed in GM1+ and GM1^−^ loci in unstimulated macrophages but formed a more stack-membrane like signal in LPS-stimulated macrophages. (G) Quantifying co-localization between GM-1 and DAT using Pearson’s correlation coefficient showed LPS-stimulated macrophages has significantly increased DAT-CTxB co-localization compared to unstimulated macrophages (p=0.0354, unpaired t-test). Images and data in F-G are from n=5, 8 experiments for unstimulated and LPS-stimulated groups, respectively.

DAT-dependent dopamine efflux is associated with an inwardly facing conformation, whereas uptake is associated with an outwardly facing conformation. This prompted us to investigate if LPS induced an inward facing conformation of DAT leading to decreased uptake, decreased inward current, and increased efflux without affecting membrane DAT levels. To quantify the levels of inward vs. outward facing conformation of DAT, we used JHC1-064, as this compound is membrane impermeable and only binds to the outwardly facing conformation of biogenic amine transporters (Lebowitz et al. 2019) (**Figure 2E**). Adopting this approach, we assessed JHC1-064/DAT binding in unstimulated and LPS-stimulated macrophages in the presence of a NET blockade using confocal microscopy (**Figure 7C**). LPS-stimulation significantly reduced the magnitude (**Figure 7D,** p<0.05) and rate (**Figure 7E**, p<0.0001) of JHC1-064/DAT binding. Indeed, the JHC1-064/DAT binding in LPS-treated macrophages was comparable to that observed in macrophages treated with nomifensine. The decreased JHC1-064/DAT binding observed in LPS-stimulated macrophages indicate that there was a significantly less outwardly facing DAT at the membrane. Collectively these data support that notion that LPS favors an inwardly facing, efflux-promoting DAT on human macrophages.

Inwardly facing and effluxing DAT has been reported to localize to GM1(Butler et al. 2015, Cremona et al. 2011) and syntaxin 1a-enriched (Binda et al. 2008) portions of the plasma membrane. Thus, we compared the co-localization of DAT and GM1 marker CTxB-555 using fixed-cell confocal microscopy (**Figure 7F**). Quantifying co-localization via Pearson’s correlation of fluorescence intensities revealed LPS stimulation significantly increased DAT-CTxB-555 co-localization (**Figure 7G,** p=0.0354). This suggests that a larger proportion of DAT localizes to GM1-enriched portions of the plasma membrane following LPS stimulation. It is important to acknowledge the dynamic nature of DAT activity—it is not an all-or-none, uptake-or-efflux dichotomy. Rather, there is a balance between DAT localizations and activities that summate to give a bulk picture, which we can observe empirically. In this context, our findings indicate that LPS-stimulation switches the dominant mode of DAT activity, favoring an inwardly facing and efflux promoting DAT preferentially localized to GM1-enriched regions of the membrane on human macrophages. Collectively, these data demonstrate a novel, dynamic mechanism by which macrophages could release dopamine in response to immune stimuli.

### CD14-dependent regulation of DAT activity engages an autocrine loop to modulate macrophage immune response

We next sought to elucidate the mechanism of (i) LPS-induced regulation of DAT activity and (ii) how DAT-dependent efflux contributed to DAT’s immunomodulatory role. On macrophages, LPS initially binds CD14 which recruits Toll-like receptor 4 (TLR4), triggering a signaling cascade that activates NF-kB and induces an inflammatory response (Płóciennikowska et al. 2015). Therefore, we hypothesized that either TLR4 or CD14 may mediate the LPS-induced regulation of DAT activity. This hypothesis was examined using three complementary approaches, an IDT307 uptake assay, DAT/GM1 localization and measurement of DAT-dependent efflux in unstimulated and LPS-stimulated macrophages. For the uptake assay, macrophages were treated with either vehicle (unstimulated), LPS, LPS + CLI095 (TLR4 antagonist, 3 µM), LPS + a neutralizing antibody against CD14 (AbCD14), or LPS + Iaxo102 (CD14 antagonist (Iaxo102, 5µM). Cells were treated for 6 hours and assayed for DAT-dependent uptake as above (**Figure 7A, B**). as pilot studies demonstrated that LPS-treatment for 6 hours produced a similar amount of DAT-dependent IDT307 uptake as did a 24-hour treatment (**Supplemental Figure 7A, B**, p=0.4183).

As expected, LPS significantly inhibited DAT-dependent IDT307 uptake [8C, unstimulated vs. LPS, p<0.0001]. Blocking CD14 with either a neutralizing antibody or small molecule antagonist abrogated this effect, returning both the magnitude and the rate of uptake to levels seen in unstimulated macrophages [**Figure 8C**, LPS vs LPS+AbCD14, AUC p<0.0001, Average slope, p=0.0636; LPS vs LPS+Iaxo102, AUC p<0.0001, Average slope LPS vs LPS + Iaxo102 p=0.0563]. The LPS-induced decrease in uptake was unaffected by TLR4 inhibition, as concurrent treatment with CLI095 did not alter the LPS-mediated decrease (**Figure 8C,** AUC vs LPS p=0.3826). The lack of effect was not due to issues with TLR4 binding by CLI095, as concurrent treatment with this compound did block the LPS induced increases in IL-6 production (**Supplemental Figure 7C**, p=0.0323).

**Figure 8:**
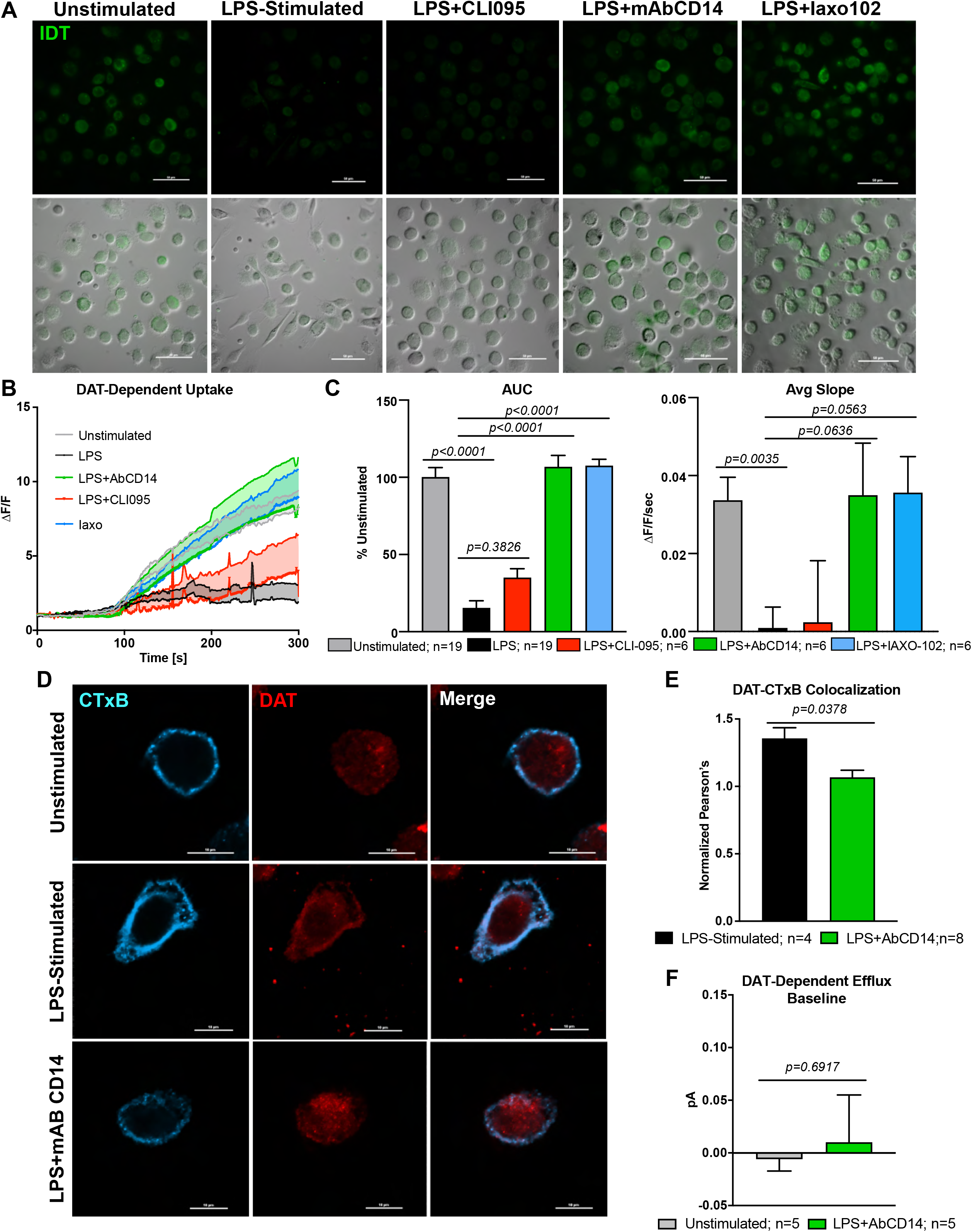
LPS-regulation of DAT activity is CD14-dependent and TLR4-independent. (A) Macrophages were unstimulated, LPS-stimulated, or co-treated with LPS and CLI095 (TLR4 antagonist), LPS and a neutralizing monoclonal antibody against CD14 (mAbCD14), or LPS and Iaxo102 (a CD14 antagonist) and assayed for DAT uptake via IDT307 in the presence of NET/OCT blockade as in Figures 3 and 6. (B) DAT-dependent uptake of IDT307 was calculated via blocker subtraction showing decreased DAT-dependent uptake in LPS-stimulated macrophages. Macrophages treated with LPS+CLI095 showed similar DAT-dependent IDT307 uptake compared to LPS-stimulated, whereas macrophages treated with LPS and either mAbCD14 or Iaxo102 showed DAT-dependent IDT307 uptake comparable to unstimulated macrophages. (C) Quantifying the magnitude (AUC, left) and rate (Average slope, right) of DAT-dependent IDT307 uptake shown in (B) showed a significant decrease in IDT307 uptake induced by LPS (AUC p<0.0001, Avg rate p=0.0035). Co-treatment with CLI095 had no significant effect on DAT-dependent uptake compared to LPS-stimulation (AUC p=0.3826), but co-treatment with either mAbCD14 or CD14 antagonist Iaxo102 rescued the DAT-dependent uptake back to unstimulated levels (AUC LPS-stimulated vs. mAbCD14 and AUC LPS-stimulated vs. Iaxo102 p<0.0001, Average slope LPS-stimulated vs mAbCD14 p=0.0636, Average slope LPS-stimulated vs Iaxo102 p=0.0563). Images and data are from n=19 experiments for unstimulated and LPS-stimulated groups and n=6 experiments/group for the inhibitor treatments. (D) Representative confocal images of fixed cultured human macrophages that were either unstimulated, LPS-stimulated, or treated with LPS and mAbCD14 labeled for GM1 (a membrane marker using CTxB-555) and DAT. LPS-stimulated macrophages exhibited more of a stacked membrane signal for DAT, which was reversed by mAbCD14. (E) Quantifying co-localization between CTxB55 and DAT using Pearson’s correlation coefficient shows that mAbCD14 reversed the increased DAT-CTxB555 colocalization back to unstimulated levels (p=0.0378, unpaired t-test). Images and data in D-E are from n=4-8 experiments/group. (F) DAT-dependent dopamine efflux measured via simultaneous whole-cell patch clamp and amperometry as in Figure 7 shows no significant increase in basal dopamine efflux between unstimulated macrophages and macrophages treated with LPS and mAbCD14 (p=0.6917, unpaired t-test, n=5 experiments/group).

Based on these results, we examined the effects of CD14 blockade on the LPS mediated increase in DAT localization to GM1-enriched areas of the plasma membrane. Analysis of CTxB-555 and DAT colocalization in vehicle, LPS and LPS + AbCD14 treated macrophages by fixed cell confocal microscopy showed that treatment with AbCD14 significantly decreased DAT-CTxB co-localization (**Figure 8D, E**, p=0.0378). This suggests that blockade of CD14 decreased the LPS-induce localization of DAT to GM1 enriched regions of the plasma membrane. Furthermore, basal DAT-dependent efflux was not different between unstimulated macrophages and macrophages treated with LPS+AbCD14 (**Figure 8F,** p=0.6917). Taken together, these data are consistent with the interpretation that LPS-regulation of DAT on macrophages is CD14-dependent.

Our findings to this point indicate that (1) DAT dynamically regulates the availability of dopamine in a macrophage’s immediate microenvironment via uptake or, during LPS-induced inflammation, via dopamine efflux (**Figures 3, 6, and 7**); and (2) DAT is an immunomodulator of the macrophage response to LPS (**Figure 5**). These findings corroborate previous research have shown that macrophages contain the requisite machinery for dopamine synthesis and dopamine signaling, the latter of which has been shown to extensively regulate macrophage immune functions (Pinoli et al. 2017). Non-neuronal cell types have recently been shown to release dopamine and signal onto their own or neighboring dopamine receptors (Aslanoglou et al. 2021). Additionally, immune cells specifically have been shown to engage in autocrine/paracrine signaling for a variety of neurotransmitters (Franco et al. 2007). Hence, a switch from DAT-mediated removal of dopamine from the extracellular milieu to DAT-mediated release of dopamine to the extracellular milieu may engage dopamine receptors to mediate DAT’s immunomodulatory effects via an autocrine or paracrine loop. To test this, macrophages were exposed to 4 distinct treatments designed to pharmacologically dissect our proposed autocrine loop (**Figure 9A**); (1) LPS treatment to increase DAT efflux and extracellular dopamine; (2) LPS treatment while blocking DAT to decrease extracellular dopamine, (3) LPS treatment while blocking DAT and adding dopamine back extracellularly which we hypothesized would reverse the effect of DAT blockade, and (4) subsequently blocking dopamine receptors which we hypothesized would resemble LPS with DAT blockade (condition 3). All conditions were tested in the presence of NET blockade. The macrophage response to each condition was then evaluated by measuring phagocytic activity, enabling evaluation of the effects of DAT-mediated dopamine release on a classical macrophage function (**Figure 9B**).

**Figure 9:**
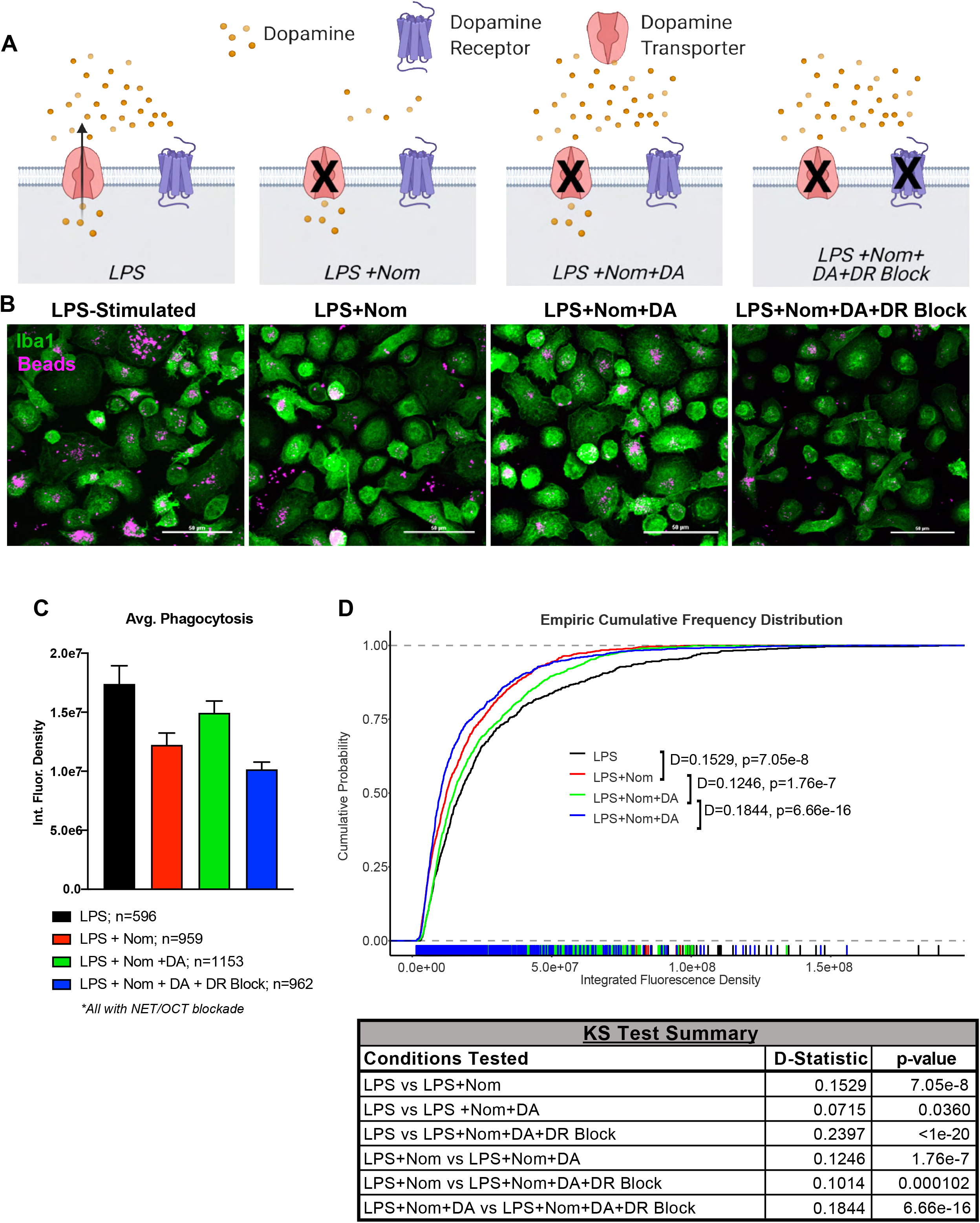
DAT immunomodulation is mediated by an autocrine/paracrine dopamine signaling loop. (A) Cultured macrophages were treated with vehicle (unstimulated), LPS to induce DAT efflux and increase extracellular dopamine, LPS+ Nomifensine to block DAT efflux and decrease extracellular dopamine, LPS+ Nomifensine+ Dopamine, LPS+ Nomifensine+ Dopamine+ Dopamine receptor blockade (Sulpiride and SCH53390) in the presence of NET/OCT blockade. (B) Representative confocal images of fixed cultured human macrophages treated with conditions as above in the presence of NET/OCT blockade and incubated with fluorescent latex beads to measure phagocytosis (unstimulated condition not shown). (C) Median phagocytic capacity measured as fluorescence intensity of phagocytic beads/cell. (D) Empiric cumulative frequency distribution curves of phagocytic capacity shows that LPS+ Nomifensine decreases phagocytosis compared to LPS-stimulated macrophages (D statistic=0.1529, p=7.05e-8). Restoring extracellular dopamine signaling by adding exogenous dopamine shifted the distribution curve to the right towards the LPS group representing an increase in phagocytosis (D-statistic =0.1246, p=1.76e-6). Blocking both D1-like and D2-like receptors reversed the effect of extracellular dopamine, shifting the distribution curve back to the left towards the LPS+ Nomifensine curve, representing a decrease in phagocytosis compared to LPS+ Nomifensine+ Dopamine (D-statistic=0.1844, p=6.66e-16). Details of all K-S tests are depicted in tabular format in (D). Images and data from (B-D) are from n=596-1153 cells/group from at least 3 experiments/group.

Quantifying the amount of phagocytosis by measuring the amount of fluorescence within each macrophage showed that blocking DAT efflux decreased phagocytic capacity compared to LPS alone (**Figure 9C, D**, D-statistic = 0.1529, p=7.05e-08), as we showed previously (**Figure 5B-E**). Macrophages from condition 3, which restored dopamine signaling by adding exogenous dopamine, showed a reversal of the effects of DAT blockade and increased phagocytic capacity approximating the LPS alone condition (LPS+Nom vs LPS+Nom+DA D-statistic=0.1246, p=1.76e-07). Macrophages from condition 4, which inhibited the exogenous dopamine signaling by blocking dopamine receptors, showed decreased macrophage phagocytosis approximating the levels seen in cells treated with LPS+Nom levels (D-statistic = 0.1844, p=6.66e-16) indicating dopamine’s effect on phagocytosis was specific to dopamine receptors. Collectively, these findings suggest that DAT efflux increases extracellular dopamine signaling to modulate macrophage phagocytosis during LPS stimulation. Altogether, our data collectively support the hypothesis that LPS stimulation of human macrophages induces a CD14-dependent shift of DAT activity favoring efflux which subsequently engages an autocrine/paracrine dopamine loop to modulate macrophage immune response.

### Parkinson’s disease macrophages exhibit a disrupted DAT-immune axis

Thus far, our data indicate that DAT activity dynamically modulates dopaminergic tone around macrophages. In this context, DAT exerts an immunomodulatory effect on the macrophage inflammatory response. We next sought to investigate if this DAT-immune axis was intact in a disease state.

Parkinson’s disease (PD) is classically characterized by loss of CNS dopamine due to death of dopaminergic neurons in the substantia nigra. Recent literature indicates that there is also an element of peripheral inflammation in PD (Harms, Ferreira and Romero-Ramos 2021, Tan et al. 2020, Dzamko, Geczy and Halliday 2015). Specifically, elevated levels of pro-inflammatory cytokines have been shown in the CNS and periphery of animal models (Sliter et al. 2018, Kozina et al. 2018, Challis et al. 2020) and PD patients (Devos et al. 2013, Reale et al. 2009, Mogi et al. 1994). Moreover, pro-inflammatory cytokines can cause dopaminergic neuron dysfunction and death and may contribute to neurodegeneration (Sommer et al. 2018, Kozina et al. 2018, Arimoto et al. 2007). Since our data indicate DAT alters macrophage cytokine secretion, we examined whether DAT activity modulated the cytokine release from PD macrophages as it did in healthy macrophages.

To this end, we treated MDM cultured from PD patients with either vehicle, nomifensine, LPS, or LPS + nomifensine and measured their release of pro-inflammatory cytokines IL-6, TNFα, and CCL2. As expected, LPS significantly increased the release of all three cytokines from PD macrophages. However, blockade of DAT during LPS stimulation had no effect on LPS induced release of IL-6 or TNFα and decreased LPS induced release of CCL2 (**Figure 10A**). This was in stark contrast to the pattern we saw in healthy macrophages (**Figure 5A**), where DAT blockade consistently increased LPS-induced release of pro-inflammatory cytokines. Comparing the effect of DAT blockade on LPS-induced release of IL-6, TNFα, and CCL2 in healthy vs. PD patient macrophages showed that PD macrophages exhibited a muted or significantly decreased response to DAT blockade (**Figure 10B**). Similarly, when we measured mitochondrial superoxide levels in PD macrophages, we noticed that DAT blockade in the presence of LPS had no effect (**Figure 10C**), in contrast to what we observed in healthy macrophages (**Figure 5F, G**). Together these data indicate that DAT-mediated immunomodulation is dysregulated in PD patient macrophages.

**Figure 10:**
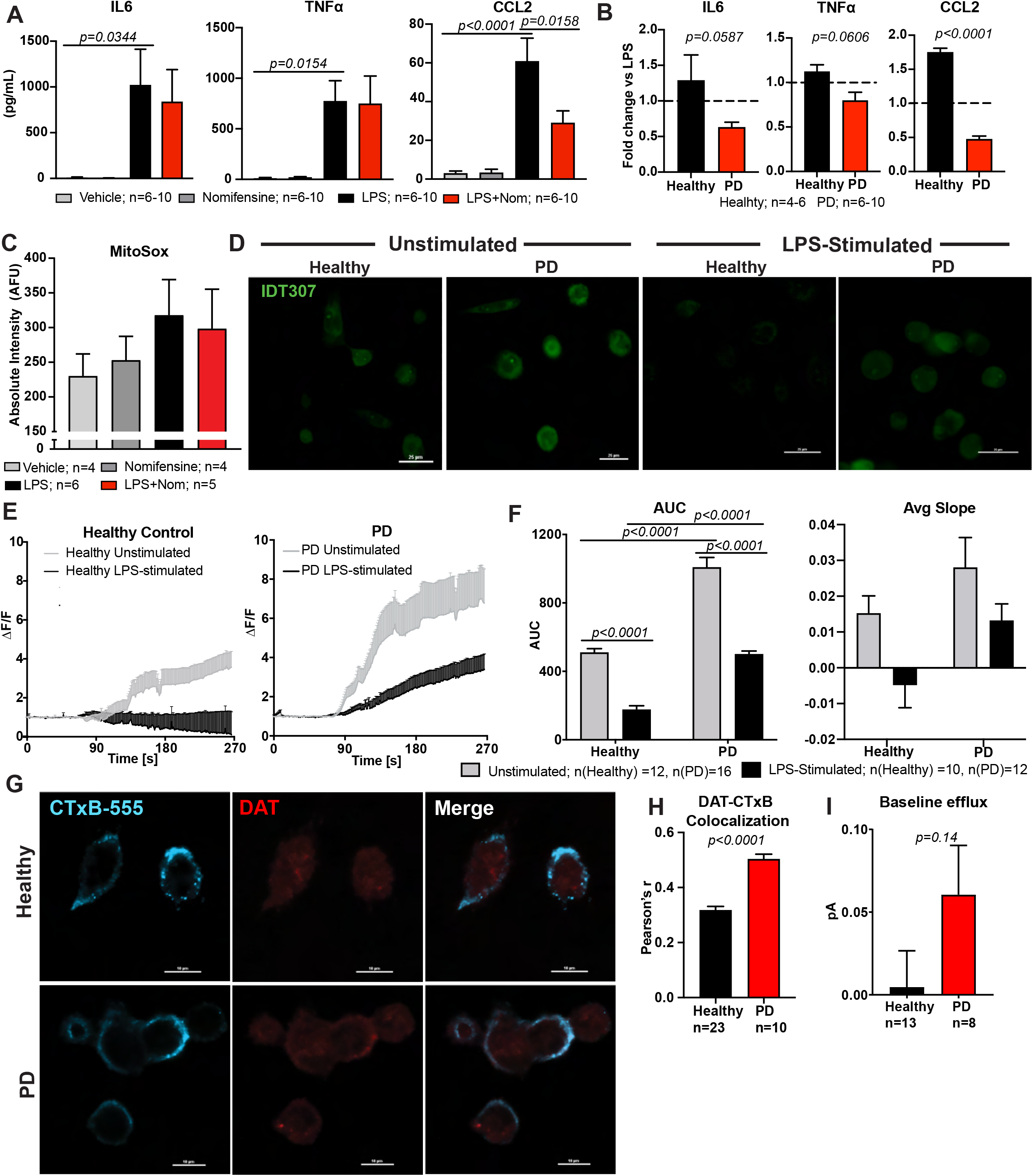
DAT immuno-modulation is disrupted in PD macrophages. (A) Macrophages cultured from PD patients (n=5-6) were treated with vehicle (unstimulated), Nomifensine, LPS, or LPS+ Nomifensine and their conditioned media was collected for cytokine analysis. LPS induced a significant increase in release of IL-6 (p=0.0344), TNFα (p=0.0154), and CCL2 (p<0.0001) from PD macrophages. Blockade of DAT in the presence of LPS had no significant effect on release of IL-6 or TNFα and significantly decreased release of CCL2 (p=0.0158). Data are from n=6-10 experiments and analyzed via one-way ANOVA with Tukey’s test for multiple comparisons. (B) The ratio LPS+ Nomifensine: LPS induced cytokine release was compared in PD macrophages vs healthy age-matched controls for IL-6, TNFα, and CCL2. The effect of DAT blockade on LPS-induced cytokine release was altered on PD macrophages (IL-6 p=0.0587, TNFα p=0.0606, CCL2 p<0.0001). Data are from n=4-10 experiments and analyzed via unpaired student’s t-test. (C) Macrophages cultured from PD patients and were treated as in (A) and assayed for mitochondrial superoxide levels using MitoSox Red™ dye under live-cell imaging. LPS+ Nomifensine did not produce a significant increase in mitochondrial superoxide levels. (D) Representative images of cultured PD patients’ macrophages and healthy age-matched controls that were treated with either vehicle (unstimulated) or LPS then assayed for DAT-mediated uptake using IDT307 under live-cell imaging as in Figures 3, 6, and 8. (E) DAT-dependent uptake was calculated for all groups using blocker subtraction as described in Figure 3 and showed that LPS induced decreased DAT-dependent IDT307 uptake in both healthy and PD macrophages. (F) Quantifying the uptake curves in (E) showed that unstimulated PD macrophages had increased magnitude (AUC, left, p<0.0001) and rate (Average slope, right) of IDT307 uptake compared to unstimulated healthy controls. While LPS significantly decreased the DAT-dependent uptake in PD macrophages (AUC p<0.0001), DAT uptake was still higher compared to LPS-stimulated, healthy controls (AUC p<0.0001). In fact, LPS-stimulated uptake in PD macrophages resembled healthy unstimulated levels. Images and data in (D-F) are from n=10-16 experiments/group, and statistics were run using a two-way ANOVA with Sidak’s multiple comparison test. (G) Representative confocal images of fixed macrophages cultured from PD patients and healthy age-matched controls labeled for membrane marker GM1 using CTxB-555 and for DAT. (H) Quantifying co-localization between CTxB-555 and DAT using Pearson’s correlation coefficient showed PD macrophages exhibited significantly increased DAT co-localization with membrane marker CTxB-555 (GM1) compared to healthy controls (p<0.0001, unpaired t-test). Images and data in (F-G) are from 10-23 experiments/group. (I) DAT-dependent dopamine efflux was measured in macrophages cultured from PD patients and healthy age-matched controls as in Figure 7. Baseline efflux was not significantly altered between PD and healthy macrophages, although there was a trended increase (p=0.14, unpaired t-test). Data are from 8-13 experiments/group.

Our data indicate that the LPS-induced switch from DAT uptake to efflux is critical to DAT-mediated immunomodulation, so we hypothesized that DAT activity and regulation were altered on PD patient macrophages leading to disrupted DAT-mediated regulation of cytokines. To assess this we compared DAT-dependent uptake of IDT307 in PD patient macrophages and healthy age-matched control macrophages at baseline and following LPS stimulation (**Figure 10D, E**). PD macrophages exhibited increased baseline uptake compared to their healthy counterparts (**Figure 10F**, AUC at baseline healthy vs. PD, p<0.0001). Moreover, PD macrophages exhibited significantly increased uptake following LPS stimulation compared to healthy, LPS-stimulated macrophages (**Figure 10F**, AUC following LPS p<0.0001). In fact, DAT-dependent uptake in PD macrophages stimulated with LPS approximated uptake seen in unstimulated, healthy macrophages. These data indicate that, in contrast to healthy macrophages that exhibit a near complete switch in the mode of DAT activity from uptake to efflux following LPS (**Figures 6 and 7**), DAT on PD macrophages has a partial response to LPS—more DAT remains in uptake mode. Supporting this idea, PD macrophages exhibited increased co-localization with membrane marker CTxB-555 (**Figure 10 F, G**) and mildly elevated basal efflux, although this did not reach statistical significance (**Figure 10H**). Overall, these findings indicate that increased DAT activity on PD macrophages leads to an incomplete response to LPS subsequently disrupting DAT-mediated cytokine modulation. Collectively, these data reflect the relevance of proper DAT regulation and our proposed DAT-immune axis on macrophages in disease.

## Discussion

Monocyte-derived macrophages are a critical population of innate immune cells with both protective and pro-inflammatory roles in a variety of tissues. Over the past two decades, an increasing amount data has suggested that biogenic monoamines, such as norepinephrine and dopamine, have immunomodulatory effects on human macrophages (Scanzano and Cosentino 2015, Pinoli et al. 2017). In addition to adrenergic and dopaminergic receptors, the norepinephrine and dopamine transporters (NET and DAT, respectively) are also expressed on macrophages (Gaskill et al. 2012, Pirzgalska et al. 2017, Mackie et al. 2018). While most research to-date has focused on receptor-mediated signaling by adding exogenous norepinephrine or dopamine, a recent study has shown that NET is a potent regulator of the phenotype of sympathetic neuron-associated adipose macrophages and is implicated in obesity (Pirzgalska et al. 2017). This study, along with the other data showing that dopaminergic systems can regulate macrophage functions, suggest that transporter biology on macrophages represents a relatively unexplored, but potentially physiologically important field. Our findings support this, presenting novel insights into how DAT activity may modulate fundamental aspects of macrophage-mediated immunity.

### The Identity of DAT+ Macrophages

In the present study, we first observed the expression of NET and DAT on circulating human monocytes, cultured monocyte-derived macrophages, and a subpopulation of intestinal macrophages in situ. In the steady state monocyte-derived macrophages contribute heavily to the macrophage pools in gut and dermis, and moderately in other tissues like lung and heart (Ginhoux and Guilliams 2016). In inflammatory states or following resident macrophage depletion, monocyte-derived macrophages will repopulate most niches (Shi and Pamer 2011). An important limitation of our cultured macrophages is that they do not entirely recapitulate the molecular identity of any macrophage *in vivo* (Gosselin et al. 2014, Amit, Winter and Jung 2016) and are mostly used to study fundamental macrophage cellular biology. Nevertheless, the use of primary human macrophages still provides a reliable human-based model system that is applicable in multiple research fields. Therefore, while we cannot say under which circumstances engrafted monocyte-derived macrophages maintain DAT expression, we found consistent DAT expression in our human model system, and it is likely that at least some monocyte-derived macrophages express DAT in either the steady state or under inflammatory conditions in tissue. This is supported by our observation that a subset of intestinal macrophages was DAT+, confirming some physiological relevance to DAT expression on macrophages. The single-cell RNA (scRNA) sequencing dataset from human colon, further confirmed that the DAT transcript was expressed in a cluster consistent with tolerizing macrophages. Notably, the scRNA-sequencing has limited resolution due to the diverse sample population (not sorted). The gut wall harbors a multitude of distinct macrophage subsets that vary based on their niche—nerve associated, blood vessel associated (De Schepper et al. 2018), lymphoid patch associated (Asano et al. 2015), and tolerizing macrophages (Hadis et al. 2011) have all been described. Our data confirm that some macrophages, mostly in the submucosa, express DAT, but these data do not elucidate the precise phenotype or ontogeny of DAT+ macrophages. As DAT+ macrophages mostly localize around MAP2+ or lymphoid follicle regions, future studies will examine more precisely the complete signature and ontogeny of DAT+ macrophages in these regions. In addition, recent advances in CyTOF and RNA-sequencing may provide more insights into DAT expression on macrophages of other tissues, identifying the specific transcriptomic identities that correlate with DAT expression.

### The role of macrophage DAT in the steady state

Our study first examined the baseline function of NET and DAT on cultured monocyte-derived macrophages and found them to be capable of uptake. As biogenic monoamine transporters provide tight regulation of monoamine tone, our data support the notion that NET and DAT on macrophages help control the monoamine concentration of present in the proximal microenvironment of each macrophage. Because these experiments were done *in vitro* in macrophage monoculture, we can say with certainty that DAT-dependent uptake is mediated by macrophages and not neurons or other cells.

Conversely, a limitation of our in vitro approach is that macrophages do not exist in isolation *in vivo* but interact heavily with their tissue niche neighbors so it is difficult to determine precisely how macrophage uptake of dopamine would affect the local environment. However, some tissue environments that are rich in macrophages, such as the gut, are also rich in dopamine. It is conceivable that the DAT-mediated uptake on macrophages may help regulate the overall dopaminergic tone of their niche. Indeed, here we show that some gut macrophages are DAT+, and such a role has already been shown for NET in adipose macrophages (Pirzgalska et al. 2017). The notion that macrophages can modulate extracellular dopamine content via the DAT-dependent uptake at rest is highly novel and represents a foundation for further investigation in multiple research fields. Such studies may use conditional and inducible knockouts specific to macrophages, e.g., CX3CR1-Cre^ER^ mice, may help expand our knowledge of how macrophage DAT contributes to baseline tissue functioning as part of a complex system. Nevertheless, the notion that macrophages can modulate extracellular dopamine content via the DAT-dependent uptake at rest represents a foundation for further investigation in multiple research field.

### Bidirectional regulation between DAT and immunity

The central findings in this study are the novel discoveries that (1) LPS-stimulation induces DAT-dependent efflux and (2) that in turn DAT activity shapes the inflammatory response to LPS stimulation. This study used LPS as an inflammatory stimulus due to its potency as an immune stimulant, and since increased LPS in the gut is associated with a variety of diseases. However, LPS is not the only stimulus that can evoke a macrophage response. Interferons, interleukins, and a host of other pathogen-associated molecular patterns (PAMPs) and damage-associated molecular patterns (DAMPs) bind to macrophage receptors (Zindel and Kubes 2020). Further complicating this picture are the intricate mechanisms regulating DAT activity by affecting trafficking (Melikian and Buckley 1999, Foster et al. 2008), multimer formation (Siciliano et al. 2018), conformational state (Hong and Amara 2010, Kniazeff et al. 2008), membrane potential (Richardson et al. 2016), and transport kinetics (Gu, Wall and Rudnick 1994, Foster and Vaughan 2011). Indeed, here we show that one classic mechanism of DAT regulation, PMA-induced internalization, is preserved in macrophages, and it is likely that others are as well. Moreover, given their different signaling pathways, it is highly probable that different immune signals have divergent effects on DAT activity. The report of one immuno-regulatory mechanism of DAT activity here provides the precedent for further investigation.

Why study DAT activity on macrophages under different paradigms? Our assessment of DAT’s ability to modulate the immune response consistently implied DAT plays an important role in down modulating the inflammatory response. Inhibiting DAT during LPS-stimulation skewed the macrophage phenotype to a more pro-inflammatory state characterized by increased cytokine production and decreased phagocytosis. This was due to the efflux of dopamine increasing local dopaminergic tone and enhancing dopamine receptor signaling. An increase in DAT-mediated dopamine release also seems consistent with a previous report showing LPS upregulates catecholamine synthesis (Flierl et al. 2007). However, it is important to consider the nuances of DAT-regulation of dopamine homeostasis. It follows that DAT’s immunomodulatory properties may also be influenced by additional mechanisms mediated by other factors, such as, for example, intracellular dopamine. Cytosolic dopamine can induce oxidative stress (Mosharov et al. 2009), which is consistent with the elevated mitochondrial superoxide species observed in our study. Indeed, mitochondrial function and metabolism are potent regulators of myeloid phenotype (Minhas et al. 2019, Minhas et al. 2021, Mills et al. 2017). Alternatively, it has been proposed that cytosolic dopamine may enter the nucleus and regulate transcription (Bergquist et al. 1998, Bergquist, Ohlsson and Tarkowski 2006). Additionally, macrophages may exhibit vesicular dopamine release, as lymphocytes exhibit a partially calcium-dependent release of norepinephrine (Musso et al. 1998), adding another variable in local dopamine homeostasis. While DAT plays a major role in the regulation of local dopaminergic tone, the network of mechanisms controlling dopamine inside and outside the macrophage are likely a composite of the above, and further study utilizing technologies like fluorescent dopamine sensors (Patriarchi et al. 2018) are needed to provide further clarity.

Irrespective of the intricacies above, our findings robustly indicate DAT’s potential to modulate immunity. These data also corroborate a previous study showing increased pro-inflammatory cytokine production from splenic macrophages in DAT−/− mice (Kavelaars et al. 2005). While these data provide a limited glimpse into macrophage functions, they raise the question of whether other macrophage functions, such as antigen presentation or chemotaxis, are affected by endogenous dopamine regulation. Indeed, dopamine receptor activation on T-cells during antigen presentation was associated with increased activation (Gonzalez et al. 2013). Moreover, it is tempting to ask how DAT function may also affect more specialized macrophage functions, such as neuronal support in the gut (De Schepper et al. 2018, Gabanyi et al. 2016). Using this study as a foundation establishing the relevance of DAT in immunity, future studies can employ co-culture systems and *in vivo* tools such as fluorescent reporters for T-cell activation (Galeano Niño et al. 2020) to further unravel further roles for the various modes of DAT activity on macrophages.

### Macrophage DAT in disease

Aberrant macrophage responses are associated with a wide variety of diseases. Lung-resident macrophages contribute to pulmonary fibrosis (Aran et al. 2019), intestinal macrophages play a role in IBD (Na et al. 2019), cardiac macrophages are associated with arrhythmias (Hulsmans et al. 2017) and cardiomyopathy (Dick et al. 2019), and infiltrating macrophages play a role in multiple sclerosis (Jordão et al. 2019). Typically, macrophages in these diseases lie to one extreme of the inflammatory spectrum—i.e., hyper-polarized towards either pro-fibrotic or pro-inflammatory phenotypes. Our data show that loss of DAT activity enhances the inflammatory behaviors of macrophages. A combination of analysis from human patient samples and mouse models (e.g., bone marrow reconstitution using DAT −/− mice into a disease model) will elucidate how macrophage DAT can be leveraged in these disease states.

Parkinson’s disease, while not considered a classic neuro-inflammatory or autoimmune disease, is associated with chronic peripheral inflammation (Tan et al. 2020, Harms et al. 2021, Dzamko et al. 2015). Elevated pro-inflammatory cytokines have been observed in CSF (King and Thomas 2017, Mogi et al. 1994), circulation (Qin et al. 2016), and gut tissue (Devos et al. 2013) of PD patients and animal models. Our current findings suggest that, in contrast to healthy macrophages, macrophages from PD patients exhibit either a muted or opposite response to DAT blockade during LPS stimulation. We attributed this to an overall increase in DAT uptake activity—it continues to remove dopamine from the extracellular space even during LPS stimulation. Hence, the dynamics of extracellular dopamine signaling around PD macrophages are disrupted thereby perturbing the autocrine/paracrine immunomodulatory loop we characterized in healthy macrophages. Importantly, we do not claim that macrophage DAT plays a direct role in PD. While it is tempting to speculate that DAT dysregulation may be tied to the chronic peripheral inflammation in PD, targeted knockouts of macrophage DAT in PD models would be needed to evaluate that possibility. More broadly, this finding suggests that disrupting DAT dynamics on macrophages has immunological consequences in a human disease context.

## Conclusion

To our knowledge, this is the first study that provides an in-depth functional characterization of both NET and DAT on human macrophages. We introduce a novel, bidirectional regulation between macrophage DAT and immunity, specifically mediated by LPS-induced dopamine efflux that enhances an autocrine dopamine loop. Critically, this indicates that DAT may differentially regulate dopamine concentrations in the proximal microenvironment in response to distinct stimuli, triggering changes in macrophage function and suggesting an active role for DAT in the macrophage immune response. In a disease setting, altered DAT activity perturbs this balance interfering with DAT-mediated immunomodulation. Ours is not the first study to suggest the importance of biogenic amine transporters on immune cells, but it is the first to examine DAT specifically. Overall, the findings of this study clearly establish and important regulatory role for DAT in the innate immune response and should serve as a call for further examination of macrophage DAT in inflammatory and autoimmune settings.

## Methods

### Solutions and Reagents

Live-cell imaging experiments and electrophysiology experiments were performed using external solution containing 130mM NaCl, 10mM HEPES, 34mM Dextrose, 1.3mM KH_2_PO_4_, 1.5mM CaCl_2_•2H_2_O, 0.5mM MgSO_4_•7H_2_O, filtered (0.22μm sterile filter), adjusted to pH 7.4 and osmolarity between 270-300mOsm. Pipettes for electrophysiology/amperometry were filled with internal solution containing, 90mM CsCl, 0mM/30mM NaCl, 0.1mM CaCl_2_•H_2_O, 2mM MgCl_2_•6H_2_O, 1.1mM EGTA, 10mM HEPES, and 30mM Dextrose. To account for potential off-target effects due to norepinephrine and serotonin transporters, all experiments were performed in the presence of 1μM desipramine (DMI, Sigma Aldrich, St. Louis, MO), and 1μM fluoxetine (Flo, Tocris, Bristol, UK). To measure DAT-dependent effects, experiments included a DAT blockade condition using 5-20μM nomifensine (Nom, Sigma Aldrich, St. Louis, MO). The reagents’ sources, solutions’ compositions, analysis software, and the equipment employed for biochemical analysis, live or fixed cell confocal microscopy imaging, TIRF microscopy, electrophysiology and amperometry are outlined in **Supplemental Tables 1, 2, and 3**.

### Human Blood Samples

This study was approved by the University of Florida’s Institutional Review Board (IRB# 201701195). To obtain macrophages of healthy donors, blood samples were purchased from LifeSouth Community Blood Center, Gainesville, FL. As outlined in the Supplemental questionnaire 1, the donors were healthy individuals aged 20-70 years-old with any sex, were not known to have any bloodborne pathogens, and were never diagnosed with a blood disease such as leukemia or bleeding disorders. None of the donors were using any medications for an infection, nor were they on any blood thinners. Whole blood samples from Parkinson’s disease patients were obtained via an IRB-approved protocol (IRB# 201701195). Following the informed consent process, patients diagnosed with Parkinson’s disease by a board-certified Movement Disorders physician and age-matched healthy controls had 20-30ml of blood drawn. Whole blood samples were then immediately taken for processing and monocyte isolation.

### Human colon tissue

Formaldehyde-fixed, paraffin-embedded samples of human colon were obtained via an IRB-approved protocol (IRB# 202002059) from the UF Center for Translational Science Institute (CTSI) Biorepository under a confidentiality agreement. The colon samples were confirmed to be histologically normal by a board-certified pathologist. Blocks of tissue were cut into 5µm sections and mounted on slides, de-paraffinized, and rehydrated. Mounted tissue samples were immersed in Na-citrate buffer (10mM Na-Citrate, 0.05% Tween-20, pH to 6.0 with 1N HCl) then subjected to 24 hours of photobleaching under a broad-spectrum LED light at 4°C to reduce autofluorescence. Following photobleaching, antigen-retrieval was performed with the samples immersed in Na-citrate buffer and steamed at 95°C for 45 minutes. Tissue samples were blocked/permeabilized using 10% normal goat serum in 0.3% Triton-X in PBS for 3 hours at room temperature followed by additional avidin biotin blocking using a kit according to manufacturer’s instructions (BioLegend, 927301). Primary immuno-staining was performed using antibodies: chicken anti-MAP2 (1:800, BioLegend, Poly28225), rabbit anti-IBA1 (1:800, Encor, RPCA-IBA1), and rat anti-DAT (1:200, Millipore, MAB369), in 5% normal goat serum 0.3% Triton-X in PBS overnight at 4°C. Slides were washed in PBS for 3x for 20 minutes each at room temperature with gentle shaking. Secondary immuno-staining was performed using: anti-chicken AlexaFluor-488+ (1:800, Invitrogen, A32931), anti-rabbit AlexaFluor-568 (1:800, Invitrogen, A-11011), and anti-rat biotin (1:200, Vector, BA-9401) for 1 hour at room temperature. Samples underwent another round of washes as above. Tertiary immuno-staining was performed using Streptavidin- 647 (1:200, Invitrogen, S-32357) for 1 hour at room temperature followed by another round was PBS washes, with the last wash going overnight. Slides were then counterstained with DAPI (1:5000, Invitrogen, D-1306) and mounted with Fluoromount Gold and allowed to dry for 1 day.

Immuno-stained colon samples were imaged using a Nikon A1 laser scanning confocal microscope (Nikon Instruments, Melville NY) and a x40 1.3 NA oil-immersion Plan-Apo objective. DAPI, MAP2, IBA1, and DAT signals were acquired via excitation with 405, 488, 561, and 640nm solid argon lasers and detected at 450/50, 525/50, 595/50, and 700/75nm, respectively. Channels were acquired in sequential series to minimize bleed-through. Z-stacks were acquired in 0.25µm steps through the sample at various anatomical locations in the colon wall and converted to maximum intensity projections. Further processing was performed in Nikon Elements NIS Analysis software (Nikon Instruments, Melville, NY).

### Macrophage Isolation and Culturing

Macrophages were isolated from human blood using Ficoll density gradient separation as previously described (Dagur and McCoy 2015). Briefly, blood was diluted with 1x PBS 1:1 ratio and then overlaid on the Ficoll for a final volume ratio of 5:5:3 for blood: PBS: Ficoll, respectively. After centrifugation, the buffy coat layer was extracted, washed with PBS, re-suspended in Monocyte Adhesion Medium (MoAM) and then plated on either 12mm #1.5 round glass cover slips or glass-bottomed dishes, and incubated for 1-2 hours at 37°C with 5% CO_2_ to allow monocytes to adhere(Butler et al. 2015). MoAM was made fresh for each culture using 7.5% autologous serum in 1640 RPMI (Corning, Corning NY) without any supplements. After the incubation, plates and dishes were washed 3x with RPMI, given fresh MoAM, and returned to the incubator. Media changes were done 1 day and 3-4 days post-culture. All experiments were run on days 6 or 7, except where explicitly stated. The experiments were performed between 4-6 days after plating.

### Flow cytometry for DAT NET and SERT

As previously published, we noted that freshly isolated PBMCs resulted in fewer than 0.5% dead cells during acquisition (Gopinath 2020); therefore, viability dye was not included in fresh PBMC samples. Every sample was stained in parallel with unstained, secondary only and isotype controls. Following PBMC isolation, cells were counted with trypan blue exclusion for viability, and density adjusted to 10,000 cells per microliter. 1 million cells per condition were aliquoted and immediately fixed for 30 minutes at room temperature (eBioscience Fix/perm kit, 88-8824-00), followed by washes in permeabilization buffer per manufacturer instructions. Primary antibodies against DAT (Sigma, MAB369, 1:100), NET (MabTechnologies, NET17-1, 1:100) and SERT (MabTechnologies, ST51-2, 1:100) were added and incubated for 30 minutes, followed by species-specific fluorochrome-conjugated secondaries (1:100 dilution). Data were acquired immediately following staining (Sony Spectral Analyzer, SP6800), and analyzed using FlowJo (Treestar).

### Flow cytometry for intracellular cytokines

Monocytes were isolated from total PBMCs prepared as described above (Gopinath et al. 2020) using negative selection (Biolegend, 480048) per manufacturer’s instructions. Total PBMCs were Fc-blocked to reduce nonspecific binding, followed by incubations with biotin-conjugated antibody cocktail containing antibodies against all subsets except CD14 (negative selection), followed by incubation with magnetic-Avidin beads, allowing all subsets other than CD14+ monocytes to be bound to the magnet. CD14+ monocytes were collected from the supernatant fraction, washed, counted and density adjusted such that 500,000 CD14+ monocytes were seeded per well (Figure SX) and treated for 6 hours with vehicle, LPS (10ng/mL), or LPS plus Nomifensine (10uM) in an ultra-low-adherence 6-well plate (Corning, 3471) to prevent adherence. Suspended cells from each treatment group were aspirated and placed in a 15mL conical tube, with any remaining adherent cells detached by incubation with 700uL Accumax solution for 3 minutes (Innovative Cell Technologies, AM105) and added to suspended cells. After pelleting cells by centrifugation (3 minutes x 100g, room temperature), cells were assayed by flow cytometry.

As previously published(Gopinath et al. 2020), cells for flow cytometry were fixed and permeabilized (eBioscience, 88-8824-00), and stained for intracellular markers MCP1/CCL2 (Biolegend, 505909, APC, 1:100), TNF-alpha (Biolegend, 502908, PE, 1:100), IL1beta (Biolegend, 511709, Pacific blue, 1:100), or IL6 (Biolegend, 501121, PE-Dazzle594, 1:100). Data were acquired immediately following staining (Sony Spectral Analyzer, SP6800), with compensation applied at the time of acquisition. Data were analyzed using FlowJo (Treestar).

### RT-qPCR

Total RNA was extracted from cells using Trizol or the RNeasy Mini Plus^™^ kit (Qiagen), and RNA quantity and purity were determined by NanoDropOne spectrophotometer (Nanodrop Technologies). RNA (1 μg) was used to synthesize cDNA from each donor using the high-capacity reverse transcriptase cDNA synthesis kit (Abcam). DAT, and the housekeeping genes UBC, OAZ1, and TBP were amplified from cDNA by quantitative PCR (qPCR) on a QuantStudio 7 using gene-specific primers. TaqMan Fast Universal Master Mix, and PCR assay probes for DAT (Hs00997374_m1), 18s (4319413E), UBC (Hs00824723_m1), OAZ1 (Hs00427923_m1), and TBP (Hs00427620_m1) genes were purchased from Applied Biosystems (ThermoFisher, Waltham, MA, USA). A ratio of the 3 housekeeping genes was calculated by taking an average which was then subtracted from the DAT value to get a normalized CT DAT value. The normalized DAT value was then transformed using the equation 2^−ΔCT^. The transformed value was then multiplied by 10,000 and plotted. Positive donors were those that expressed DAT in at least 1 of 3 wells tested.

### LPS Stimulation and Blockades

Ultra-pure lipopolysaccharide from *E. coli* was added to a final concentration of 10ng/ml. Cells were cultured as described above, supplemented with 7.5% autologous serum, then either treated with fresh media (un-stimulated) or LPS-supplemented media (LPS-stimulated) and incubated at 37°C with 5% CO_2_ for either 6 or 24 hours before beginning experiments. For mechanistic studies (Figure 8), the macrophages were pre-treated with 3μM CLI-095, a TLR4 antagonist(Ii et al. 2006) (Invitrogen, San Diego CA) for at least 1 hour (6hr LPS stimulation) or 3 hours (24hr stimulation). After pre-treatment, macrophages were treated with LPS±CLI-095 (Invitrogen, San Diego CA) at their respective concentrations for either 6 or 24 hours. Neutralizing CD14 antibody (10μg/ml, BioLegend) or a small molecule CD14 inhibitor, Iaxo-102 (5µM, AdiopoGen) was used in separate sets of experiments to test CD14-dependency of the measured responses. Macrophages were pre-treated with CD14 antibody for at least 1 hour(Kurt-Jones et al. 2000) before being treated with LPS±CD14 antibody following the same LPS-stimulation paradigm described above. For inhibitor alone control, the un-stimulated macrophages received antibody or CLI-095.

### Biotinylation and Immunoblotting

Unstimulated and LPS-stimulated macrophages were washed three times with cold PBS, then incubated with sulfo-NHS-biotin (1.5mg/mL; ThermoFisher Pierce, 21331) for 30 minutes at 4°C while rocking. Remaining sulfo-NHS-biotin was quenched with cold Quenching Solution (glycine 50mM in PBS), followed by three washes with cold PBS. Cells were lysed in BufferD lysis buffer (10% glycerol, 125mM NaCl, 1mM EDTA, 1mM EGTA, pH 7.6) containing 1% TritonX and protease inhibitor cocktail (Millipore, 539131) for 1 hour at 4°C while rocking, followed by centrifugation for 15 minutes at 12,000g. The supernatants were divided into three portions: 25uL for protein quantification, 300uL for incubation with Avidin, with the remainder for whole lysate input. After equilibrating monomeric UltraLink Avidin (ThermoFisher Pierce, 53146) twice with 1mL BufferD, 80μL of 50% bead slurry were added to 300μL lysate and incubated at 4 degrees C for 1 hour with rotation. Supernatant was retained as cytoplasmic fraction, and beads were washed three times with 1mL BufferD, eluted with 40μL Laemmli Sample Buffer 4x (containing 10% beta-mercaptoethanol) at 37°C for 30 minutes, separated by 10% SDS-PAGE, transferred to 0.45um nitrocellulose, and probed with anti-DAT antibody (Millipore-Sigma, AB5802). Fluorescent images were analyzed using Fiji/ImageJ (NIH) to measure image density of DAT bands. Values were normalized to total protein per lane. Beta-tubulin (Encor, MCA 1B12) was probed to demonstrate membrane fraction isolation during biotinylation.

### Immunocytochemistry

Un-stimulated and LPS-stimulated cells were washed 3 times with 1x external solution and then incubated with CTxB conjugated to AlexaFluor 555 (1:500, Invitrogen, C-34776) +/− LPS in 1x external solution with rotation for 30 minutes at 4°C to limit internalization. Following this, cells were fixed in 4% PFA for 30 minutes at room temperature, and then blocked/permeabilized with 5% normal goat serum, 0.5% Triton-X in PBS for 1 hour at 37°C and 5% CO_2_. After blocking, cells were stained with anti-DAT antibody (1:1000, MAB369, Millipore, Berlington MA) in 0.1% Triton-X, 5% normal goat serum in 1xPBS and incubated overnight at 4°C with rotation. Cells were then washed 3 times with PBS for 30 minutes, stained with anti-rat AlexaFluor-647 (1:1000, Invitrogen, A-21247) secondary antibody in 0.1% Triton-X, 5% normal goat serum in 1x PBS for 1 hour at room temperature, then washed again as described above. For DAT/IBA1, NET/IBA1, or SERT/IBA1 immunofluorescent labeling, this process was repeated without the initial CTxB-555 incubation. Primary antibodies, in addition to MAB369, were rabbit anti-IBA1 (1:1000, Encor, RPCA-IBA1), mouse anti-NET (1:100), and mouse anti-SERT (1:2500). Secondary antibodies included anti-rat AlexaFluor-647 (1:1000, Invitrogen, A21247), anti-mouse AlexaFluor-647 (1:1000, Invitrogen, A-21235), and anti-rabbit AlexaFluor-568 (1:1000, Invitrogen, A-11011). Cells were stored in foil at 4°C until ready for imaging. To determine specificity, one dish per experiment was stained only with secondary antibody. **Supplemental Table 2** outlines the source and concentrations of antibodies used in this study.

### Fixed Cell Confocal Imaging and analysis for DAT/NET/SERT expression and co-localization

Immuno-stained samples were imaged using a Nikon A1 laser-scanning confocal microscope (Nikon Instruments, Melville NY) using a x60 1.4NA oil immersion objective. Excitation of fluorophores was accomplished using argon lasers of wavelength 561nm and 640nm. Emission was detected at 595/50nm and 700/75nm, respectively. To prevent bleed through, signals were acquired in sequential series. Z-stack images were taken from the basal membrane plane through the top of the cell. For co-expression of IBA1 with DAT, NET, or SERT, maximum intensity projections were generated in Nikon NIS Elements Analysis software (Nikon Instruments, Melville, NY).

To calculate co-localization, for each z-stack, only the z-planes of stacked membrane were taken for analysis. Regions of interest were manually drawn around cells and co-localization between the CTxB555 and DAT signals, as measured by Pearson’s correlation coefficient, was taken as the average for each region of interest through the selected z-planes using Nikon Element NIS Analysis software. Individual Pearson’s correlation coefficient values were averaged to establish the mean Pearson’s correlation coefficient for the image. Image processing for representative images (e.g., rolling-ball background subtraction or denoising) was performed using Nikon Elements NIS analysis software (Nikon Instruments, Melville, NY).

### Fixed-cell Total Internal Reflective Fluorescence Microscopy (TIRF-M)

CFP-DAT expressing CHO cells were fixed and immuno-labelled for DAT (1:1000, MAB369, Millipore) as above. Macrophages were labelled with CTxB-555 and with anti-DAT antibody (1:1000, MAB369, Millipore), mouse anti-NET (1:100), or mouse anti-SERT (1:2500) as above. The exception is that secondary labeling was accomplished using anti-rat DyLight 405 (1:5000, Novus, NBPI-72983) or anti-mouse DyLight 405 (1:1000, BioLegend, Poly-24091). Fixed cells were then imaged on a Nikon Eclipse TE2000 inverted microscope using a 60x 1.49NA oil-immersion TIRF objective and 405nm and 560nm solid lasers using a CoolSNAP HQ^2^ CCD camera. The lasers were each aligned manually. Cells were first imaged in brightfield. The TIRF plane was identified using the CFP laser line for CFP-DAT CHO cells and the RFP laser line for macrophages. The TIRF laser incident angle was adjusted manually for each experiment. For each field of view, images in the CFP channel and RFP channel were acquired sequentially, and this process was repeated for multiple fields of view. Image processing for representative images (e.g. background correction and denoising) was performed using Nikon Elements NIS Analysis software.

### STED Microscopy

Macrophages were prepared for fixed cell imaging as described above. STED imaging was performed on an Abberior Instruments Expert Line microscope (Abberior Instruments, Germany) mounted on Olympus IX83 microscope operated by Imspector software (version 16.3.11961-w2026; Abberior Instruments, Germany). Alexa 647 was imaged with a 640-nm pulsed excitation laser and a pulsed 775-nm STED laser (5 % of 1.25 W power). Alexa 555 was imaged with a 561-nm pulsed excitation laser and the 775-nm STED laser (40 % of the power). An oil-immersion objective lens (UPlanSApo 100xO NA1.4, Olympus, Japan) was used for imaging. The xy images were recorded with a pixel size of 20 nm in the x and y directions and a pixel dwell time of 10 µs. The xz images were acquired with 100 % of 3D-STED, a pixel size of 86 nm in the z direction, and a y step size of 1 µm.

### Analysis of previously published single-cell RNA sequencing data

A unique molecular identifier count matrix was obtained from Li et al. 2020 via the NCBI’s Gene Expression Omnibus by searching for terms “human” and “gut-resident macrophage” or “human” and “intestinal macrophage”. Details regarding the acquisition of the original data set can be found in (Li et al. 2020).The resulting count matrix was imported into R using the Read10x function (part of the Seurat package). Quality control, normalization and scaling, principal component identification, clustering, and dimensionality reduction were all carried out using the Seurat pipeline, with default settings as previously described, to generate a t-SNE plot with 13 distinct clusters. Additionally, expression of PTRPP (CD45) as a feature plot to identify leukocytes, and expressions of SLC6A3, AIF1, FGCR2A, CD80, Siglec, and IL-10 as violin plots to annotate cluster 7 were all accomplished via the Seurat pipeline.

### 3H-Dopamine (^3^H-DA) Uptake

Human macrophages were plated in a 24 multi-well plate and cultured for 6 days. The experiments were performed in either external solution containing NET and OCT blockers (1μM desipramine and 1μM fluoxetine), or external solution containing NET, OCT, and DAT blockers (10μM desipramine, 1μM fluoxetine, and 5μM nomifensine). Media was aspirated and replaced with 200μL uptake buffer with either 10μM desipramine and 1μM fluoxetine or 10μM desipramine and 1μM fluoxetine and 5μM nomifensine, and 5μL of 0, 0.3, 3.0, 30, 100μM, or 300μM cold DA, then 50μL of ^3^H-DA was added to each well. The 300μM cold DA condition measures non-specific DA uptake. After incubation, the uptake solution was removed; the cells were washed, then lysed with SDS for 1 hour while rocking at room temperature. The lysate was collected, and ^3^H-DA content was measured.

### IDT307 Uptake

Human macrophages were cultured on round coverslips for 4-6 days. On the day of experiment, the coverslips were placed into an imaging chamber (Warner Instruments, RC-26G) and washed with 1x external solution. Perfusion throughout the experiments was accomplished using a Warner VC-8 automated perfusion system (Warner Instruments, RC-26G) with the flow rate adjusted to approximately 2-3ml/min. All experiments were performed on a Nikon FN-1 upright microscope (Nikon Instruments, Melville, NY) using a Nikon 40x 0.8NA NIR APO objective. A phase contrast image was taken prior to and following each experiment to monitor cell health. Throughout the experiment, a SpectraX Light Engine (Lumencor, Beaverton OR) was used to emit a 480nm excitation laser at 5% power passed through a quadruple bandpass filter (Chroma Technology Corp., Bellows Falls VT). Images were acquired every 1 second with 100ms exposure with an Andor Zyla 4.2 PLUS camera (Andor Technology, Belfast, UK).

For Figure 3, coverslips baseline and IDT perfusion solutions either contained (1) no antagonists, (2) blockade against NET/OCT (desipramine 10µM, fluoxetine 1µM) to isolate DAT; (3) blockade against DAT/OCT (nomifensine 5µM, fluoxetine 1µM) to isolate NET; and triple antagonist cocktail to eliminate specific blinding. For remaining experiments, coverslips were first perfused for 1 minute with external solution, followed by external solution containing specific NET and OCT blockers (1μM desipramine and 1μM fluoxetine), or external solution containing NET, OCT, and DAT blockers (1μM desipramine, 1μM fluoxetine, and 5μM nomifensine) to establish baseline fluorescence values. Once baseline was established, coverslips were perfused for 4 minutes with 5μM IDT307 (Sigma Aldrich, St. Louis, Mo), a fluorescent substrate for neurotransmitter transporters. In parallel experiments, the DAT-mediated substrate (IDT307) uptake was measured in the unstimulated and LPS-stimulated human macrophages. For PMA-induced change in uptake, unstimulated macrophages were treated with 1μM PMA in 1x external for 15 minutes prior to perfusion. Changes in intracellular fluorescence levels (indication of DAT-mediated substrate uptake) were measured for each cell using Nikon NIS-Elements software with background subtraction correction. Each cell served at its own control, where the fluorescence signals were normalized to the cell’s baseline fluorescence signal (average fluorescence value across first 60 seconds). DAT-dependent (nomifensine-sensitive) uptake was measured by subtracting the normalized fluorescence profile of the nomifensine blockade condition from the Desipramine/Fluoxetine condition. Area under curve for each condition was calculated using Prism 7.0 software (GraphPad) to measure cumulative uptake throughout the perfusion period.

### JHC1-064 Binding

JHC1-064 was a generous gift from Dr. Amy Newman at the National Institute for Drug Abuse. For Figure 2, macrophages were pre-treated with (1) no antagonists; (2) blockade against NET/OCT (desipramine 10µM, fluoxetine 1µM) to isolate DAT; (3) blockade against DAT/OCT (nomifensine 5µM, fluoxetine 1µM) to isolate NET; and triple antagonist cocktail to eliminate specific blinding. For Supplemental Figure 2, CFP-DAT CHO cells (positive control group) were pre-treated with either vehicle or nomifensine (5µM). For Figure 7, un-stimulated or LPS-stimulated human macrophages were incubated in the external solution supplemented with 1μM desipramine and 1μM fluoxetine for 5 minutes. The fluorescence baseline was taken for 1 minute. JHC1-064, a fluorescent analog of cocaine, which binds to DAT (Cha et al. 2005), was then added to a final concentration of 50nM. Cells were imaged on a Nikon A1 laser-scanning confocal microscope with a 60x 1.4NA Nikon Plan-Apo objective, and Nikon “perfect focus” were used for all imaging. JHC1-064 was excited using a solid argon 568nm laser. All imaging was done at 37°C. Images were analyzed by measuring the fluorescence signal in the regions of interest (ROI) manually drawn around cells. Changes in the fluorescence signal at the surface membrane after drug application or vehicle were measured and normalized to each region’s baseline fluorescence value to determine fold change in JHC1-064 binding to DAT. JHC1-064 rate of binding calculated as the average slope beginning at 60 seconds, when JHC1-064 perfusion began through the end of recording (GraphPad).

### JHC1-064 Total Internal Reflection Fluorescent (TIRF) Microscopy

For Figure 2 and Supplemental Figure 2, YFP-DAT cells (positive control group) or human macrophages were treated as above. Following binding, observed via confocal microscopy, dishes were transferred to an inverted Nikon Eclipse TE200 microscope equipped with solid 405, 515, and 560nm lasers and a 60x 1.49NA oil-immersion objective for TIRF-M imaging, as above, using the 560nm (RFP laser line only). Snapshot images of the TIRF plane were taken using the 560nm laser. For Supplemental Figure 6, macrophages were incubated in 40nM JHC1-064 solution for 30 min at room temperature as described previously(Richardson et al. 2016). The live-cell TIRF imaging was done at 37°C using an objective heater (20/20 Technology Inc). The JHC1-064 binding was confirmed via live cell imaging; dishes were then washed to remove unbound JHC1-064, then placed back on the TIRF microscopy system(Richardson et al. 2016). Baseline fluorescence was recorded for 30s before application of either vehicle or 1μM phorbol 12-myristate 13-acetate (PMA) (Sigma Aldrich, Cat# P8139) and imaged every 10s for 5 minutes at 50% laser power. The data were analyzed as we have described previously(Richardson et al. 2016). Regions of interest were manually drawn around cells and fluorescence was measured throughout the experiment in each ROI using Nikon Elements NIS analysis software. The change in fluorescence was calculated as (*f-f_baseline_*)/*f_baseline_* × *100%*.

### FFN200 Loading and Release

Un-stimulated and LPS-stimulated macrophages cultured as above were washed once with external solution and then incubated in 20μM False Fluorescent Neurotransmitter 200 (FFN200, Tocris Biosciences, Bristol UK) for 45 minutes at 37°C in the dark. Following incubation macrophages were washed once more with external and imaged. Imaging studies were all conducted at 37°C on a Nikon FN-1 upright microscope and x40 objective as described above for the IDT assays. Perfusion system is described above. A fluorescence baseline was taken for 30 seconds using a 380nm LED laser from a SpectraX Light Engine imaging once every 3 seconds. The macrophages were either perfused with vehicle (external solution containing with NET/OCT blockade), 10μM amphetamine, or 10μM amphetamine + 5μM nomifensine for 4 minutes using the same imaging settings as above. For analysis, regions of interest (ROIs) were manually drawn around cells. Fluorescence intensities at baseline and throughout the perfusion were quantified using Nikon Elements software. For baseline fluorescence, the fluorescence values measured during baseline was averaged. For the response to drug, cells’ fluorescence values were normalized to their respective baseline and multiplied by 100 yielding normalized fluorescence.

### Whole-cell Electrophysiology Recordings

Cells/macrophages were visualized using an Andor Zyla 4.2 PLUS camera and a 40x Nikon 0.8NA Plan-APO objective for the duration of the experiment on a Nikon FN-1 upright microscope in the presence of NET/OCT blockade (10µM desipramine, 1µM fluoxetine). Whole cell voltage-clamp recordings were taken using an Axopatch 200B amplifier and Digidata 1440A low-noise data acquisition system. All parameters were monitored using Axon Clampex software (Molecular Devices, Sunnyvale CA). Low noise borosilicate glass pipettes (outer diameter: 1.0mm; inner diameter: 0.78mm; Sutter Instruments Cat. BF100-78-10) were pulled using a laser-based pipette puller (Model P-2000, Sutter Instruments Co, Novato CA) with a tip resistance of 2-5MΩ. The data were analyzed using Clamp-fit software (Molecular Devices, Sunnyvale CA). First, cells/macrophages were patched in cell-attached configuration with a membrane resistance of ≥1GΩ, and then the intracellular compartment was accessed via rapid application of negative pressure, switching to whole-cell configuration. The DAT-mediated inward current was measured in YFP-DAT cells (positive control group), and in un-stimulated and LPS-stimulated macrophages via whole cell voltage-clamp recordings as we have described previously(Richardson et al. 2016). Briefly, once a stable patch-clamp was established in the whole-cell configuration at a holding voltage of -40mV, a voltage step protocol was run from 0 to -120 mV in 20mV increments to establish a baseline voltage current relationship, I(V). Each I(V) protocol was separated by at least 2-3 min to ensure cell viability. After acquiring baseline I(V), nomifensine was applied to measure uncoupled DAT-mediated current (baseline current minus current after nomifensine,10μM)(Richardson et al. 2016). The amphetamine-induced, DAT-mediated current is defined as amphetamine-induced current minus nomifensine-induced current(Richardson et al. 2016).

### Amperometry

Single cell simultaneous patch-clamp and amperometry experiments were performed as described previously(Sambo et al. 2017, Butler et al. 2015). The amperometric carbon-fiber electrode (ProCFE, Dagan Corp.) was connected to an Axopatch 200B amplifier (Molecular Devices, Sunnyvale CA). In the presence of NET/OCT blockade (10µM desipramine, 1µM fluoxetine), macrophages were patch-clamped in cell-attached configuration, first with minimum 1GΩ membrane resistance, and then switched to whole-cell configuration by rapid application of negative pressure and dialyzed with dopamine via the patch electrode(Richardson et al. 2016). The amperometry electrode was placed near the cell and held at +700mV. Oxidative currents were generated by a voltage-step protocol starting at 100mV and decreasing to 60mV in 20mV increments and recorded via the amperometry electrode. Currents were recorded at baseline (5 minutes after whole-cell configuration was achieved), 3 min post amphetamine addition, and 3 min post nomifensine addition. Recordings were low-pass filtered at 60Hz for subsequent analysis. Efflux was reported as the averaged difference between the last 100ms of the voltage step and the 100ms prior to the voltage step. Baseline and the amphetamine-induced, DAT-dependent dopamine efflux were calculated by subtracting the efflux after nomifensine treatment.

### MitoSOX™ Imaging

Human monocyte-derived macrophages from healthy donors were cultured as described above on 12 mm #1.5 round coverslips. Complete, fresh MoAM was supplied to the macrophages on day 3 and experiments were conducted on days 6-7. Macrophages were given fresh, nomifensine-supplemented, LPS-supplemented, or nomifensine- and LPS-supplemented complete MoAM and incubated at 37°C and 5% CO_2_ for 24 hours before beginning experiments.

Thirty minutes prior to live-cell imaging, 500µL of conditioned media was collected from each 1mL well for AlphaLISA assay and macrophages were loaded with MitoSOX™ Red (Sigma Aldrich, St. Louis, Mo) mitochondrial superoxide indicator at a concentration of 1µM in complete, fresh MoAM at 37°C. Coverslips were then placed in an imaging chamber (Warner Instruments) and briefly perfused with 1x external solution at room temperature prior to imaging. Experiments were carried out on the Nikon FN-1 microscope used for IDT307 uptake with the same objective lens. A 555nm excitation laser passed through a quadruple bandpass filter at 25% power during the experiment. Images were captured with an Andor Zyla 4.2 PLUS camera and analyzed using Nikon NIS-Elements software. Objects were identified based on a fluorescence intensity threshold and converted to ROIs. Mean fluorescence intensity of each ROI was taken as a measure of mitochondrial oxidation for each individual macrophage.

### Fixed-cell phagocytosis assay

We adapted a recently published protocol for fixed-cell analysis of phagocytosis *in vitro* (Caponegro et al. 2020). Cultured macrophages were treated with either (1) vehicle (complete media), (2) LPS (10ng/ml), (3), LPS + nomifensine (10µM), (4) LPS + nomifensine + dopamine (1µM), or (5) LPS + nomifensine + dopamine + Sulpiride (5µM) + SCH23390 (5µM). All conditions also contained .007g/ml of ascorbic acid to minimize dopamine oxidation (pH balanced to approximately 7.4 with NaOH) and desipramine (10µM). 1-2 wells/condition also received 0.1µg/ml of fluorescent latex beads. Macrophages were incubated at 37°C for 2hr. Following incubation, 400µl of conditioned media from wells without phagocytic beads was collected for AlphaLisa assay (see below). Macrophages were washed 3x with PBS and then fixed in 4% PFA at room temperature for 20 minutes. Following fixation, macrophages were blocked/permeabilized using 0.3% Triton-X and 5% normal goat serum in PBS for at least 1hr at 37°C. Macrophages were then incubated with anti-IBA1 from rabbit (1:1000, Encor) overnight at 4°C on a rocker. Following primary incubation, macrophages were washed 3x with PBS with moderate rocking at room temperature followed by a 1hr incubation with anti-rabbit AlexaFluor-488 (1:1000, Invitrogen) at room temperature. After 3x 20min PBS washes, coverslips were mounted and allowed to dry overnight.

Macrophages were imaged using a Nikon A1 laser scanning confocal microscope and a 40x oil-immersion 1.3NA Plan-Apo objective. The IBA1 and fluorescent beads were excited by 488nm and 561nm solid argon lasers, respectively, and detected at 525nm and 595nm, respectively. To prevent bleed through, lasers were fired in a sequential series. Multiple z-stacks taken through the depth of the cells were acquired for each coverslip. Images were then imported into Fiji/ImageJ for blinded analysis and the channels were separated. The 488nm channel (IBA1) was collapsed into a maximum intensity projection and regions of interest were manually drawn around each cell. The integrated fluorescence density for each region of interest from the 560nm channel (beads) was extracted and pasted into an excel sheet. This process was repeated for each image, from each coverslip, for each experiment. The vehicle, LPS, and LPS+Nom conditions from the first three experiments were analyzed to generate the data in Figure 5. The later experiments were analyzed to generate the data in Figure 9. The data table was uploaded into a publicly available pipeline for statistical analysis. The graphical interface generates frequency histograms and cumulative probability distributions for the data set and runs sequential Kolmogorov-Smirnov tests. These are the statistics reported in the results, figures, and figure legends. The complete code and link to the graphical user interface are available at: Simultaneously, the data were uploaded into R to generate cumulative frequency distributions aesthetically tailored to the color scheme of the rest of the figures, and the data were uploaded into GraphPad to generate plots comparing median fluorescence.

### AlphaLISA assay

Monocyte-derived macrophages were treated as described above and supernatants were collected prior to microscopy assays. Supernatants were shipped on dry ice to the Gaskill Lab at Drexel University, where they were analyzed for IL-6, TNF-α, CCL2 using AlphaLISAs performed according to the manufacturer’s protocol (PerkinElmer). Briefly, supernatants were thawed and aliquoted in duplicate into 1/2 area, opaque white 96 well plates (#6002290, Perkin Elmer) at 5 µL per well. Following addition of sample, master mix containing acceptor beads and antibody is added to each well, and plates are incubated in the dark at room temp on a high-speed shaker for 60 minutes. After 60 minutes, donor beads are added, and plate is again incubated at room temperature in the dark on a high-speed shaker for 30 minutes. After second incubation, plates are read on an Enspire Multimode Plate reader using the AlphaLISA setting. The lower limits of detection for AlphaLISAs were IL-6, 1.3 pg/ mL; TNF-α, 2.2 pg/mL; CCL2, 3.8 pg/ml. The limit of quantification for these assays was 10 pg/mL for TNF-a and IL-6, and 30 pg/mL for CCL2.

### IL-1β ELISA

Human macrophages were cultured at a density of 100,000 cells/well and treated with either vehicle, LPS, nomifensine, or LPS+nomifensine as described above. Following treatment, cells were lysed in the presence of a protease inhibitor cocktail. Cell lysate was collected, then analyzed for IL-1β using an IL-1β ELISA (BioLegend, 437004, San Diego CA) according to the manufacturer protocol.

### Statistical Analysis

Data analysis was performed via Prism 7 (GraphPad Software Inc.) or, for phagocytosis analysis, in R/an R-encoded interface. Sample sizes were not determined by sample size calculations but represent sample sizes similar to those generally used in the field for the respective types of experiments. Statistical analyses were designed assuming normal distribution and similar variance among groups tested. N was reported as number of cells when validating pharmacology and basic transporter function. Also, for phagocytosis, n was reported as number of cells to take a whole-population approach as previously described. Otherwise, n represent independent experiments. One-way, two-way, or repeated measures ANOVA (with *multiple comparison* tests as indicated for each experiment) were used for the statistical comparison of more than two means of data with a normal distribution. Unpaired t-tests were used to compare two means of data with assumed normal distributions. Kolmogorov-Smirnov tests were used to compare non-normal cumulative frequency distributions. Significance at p < 0.05 was considered statistically significant

---

**Supplemental Figure 1:**
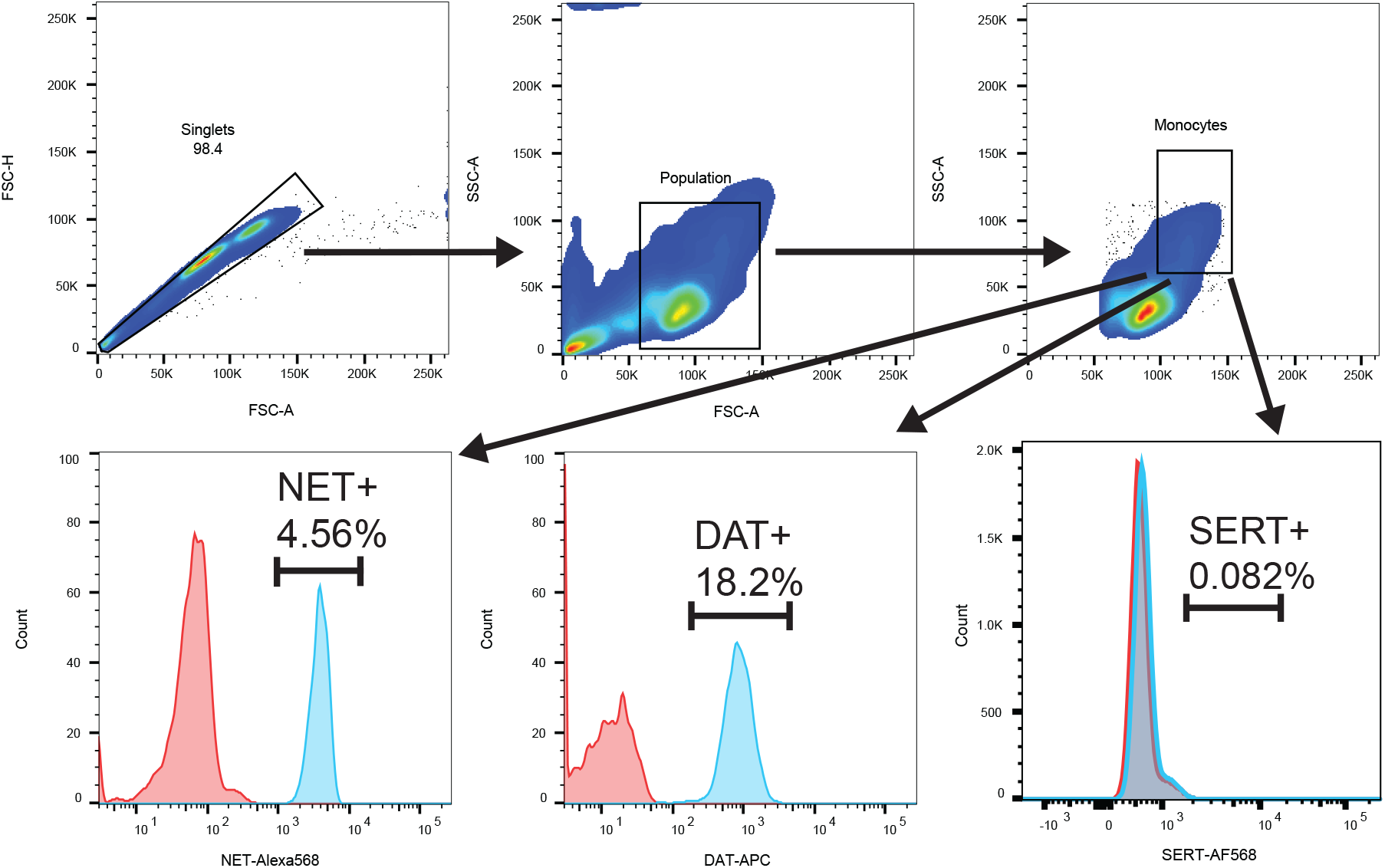
Gating scheme for flow cytometry in Figure 1. Whole peripheral blood mononuclear cells were isolated from human blood and assayed for NET and DAT expression via flow cytometry. Doublets were excluded on the basis of FSC-H and FSC-W. Monocytes were identified from the singlet population on the basis of FSC-A and SSC-A. NET- and DAT-expressing monocytes were identified from the monocyte population on the basis of NET-Alexa568 fluorescence or DAT-APC fluorescence, respectively.

**Supplemental Figure 2:**
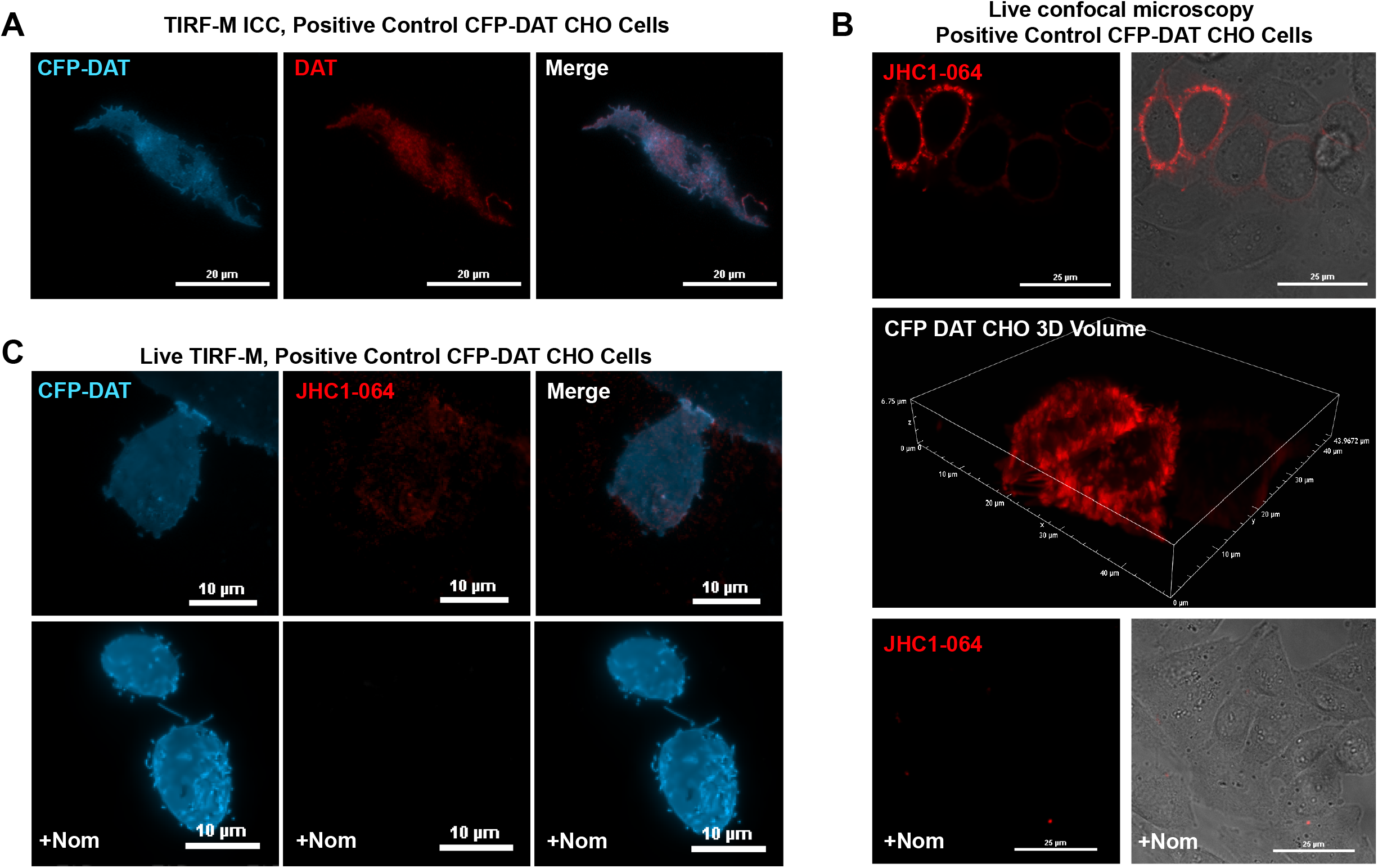
Validation of assays for detection of NET and DAT. (A) Representative images of CFP-tagged-DAT-expressing CHO cells (CFP-DAT CHO) were fixed, labelled with an anti-DAT antibody, and imaged using TIRF microscopy, showing overlap between the CFP tag and the DAT antibody labelling. (B) Representative images of CFP-DAT CHO cells incubated with JHC1-064 and monitored for binding under live-cell confocal microscopy in the absence (top) or presence (bottom) of DAT blockade (Nomifensine). No JHC1-064 fluorescence was observed when DAT was blocked. (C) Representative TIRF-M images of CFP-DAT CHO cells following JHC1-064 binding in the absence (top) or presence (bottom) of DAT blockade (Nomifensine). No JHC1-064 fluorescence was observed on CFP+ cells in the presence of DAT blockade (+nomifensine). Images are from at least 3 experiments/group.

**Supplemental Figure 3:**
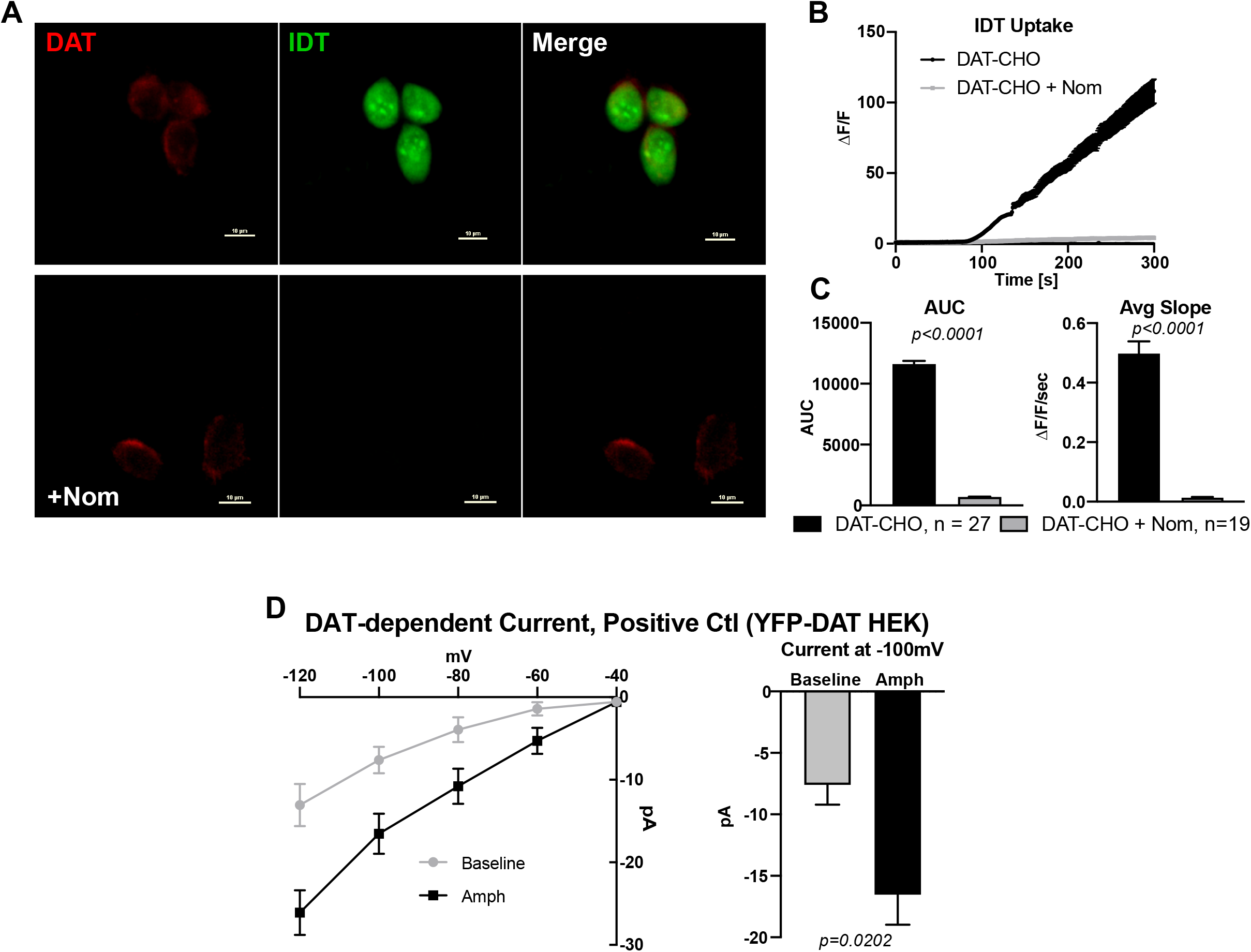
Validation of assays for functional characterization of NET and DAT. (A) Representative images of CFP-DAT CHO cells assayed for DAT-mediated uptake using IDT307 perfusion in the absence (top) or presence (bottom) of DAT blockade (Nomifensine). (B) The DAT-mediated IDT307 uptake measured as change in intracellular fluorescence above baseline (ΔF/F), revealing appreciable IDT307 uptake in the absence of DAT-blockade. (C) Nomifensine significantly inhibited the magnitude (AUC, left, p<0.0001, unpaired t-test) and rate (Average Slope, right, p<0.0001, unpaired t-test) of DAT-mediated IDT703 uptake, indicating DAT-specificity. Images and data are from 19-27 CFP+ cells over 3 experiments. (D) HEK cells stably expressed YFP-tagged DAT (YFP-DAT HEK cells) were patch clamped in whole cell configuration under voltage clamp mode, and inward currents were evoked by serial hyperpolarization steps. DAT-dependent current was calculated by subtracting the inward currents in the presence of DAT blockade from the inward currents at baseline or in the presence of DAT-activator amphetamine (Amph). Amphetamine significantly increased the DAT-dependent inward current (p=0.0202, unpaired t-test, n=5-8)

**Supplemental Figure 4:**
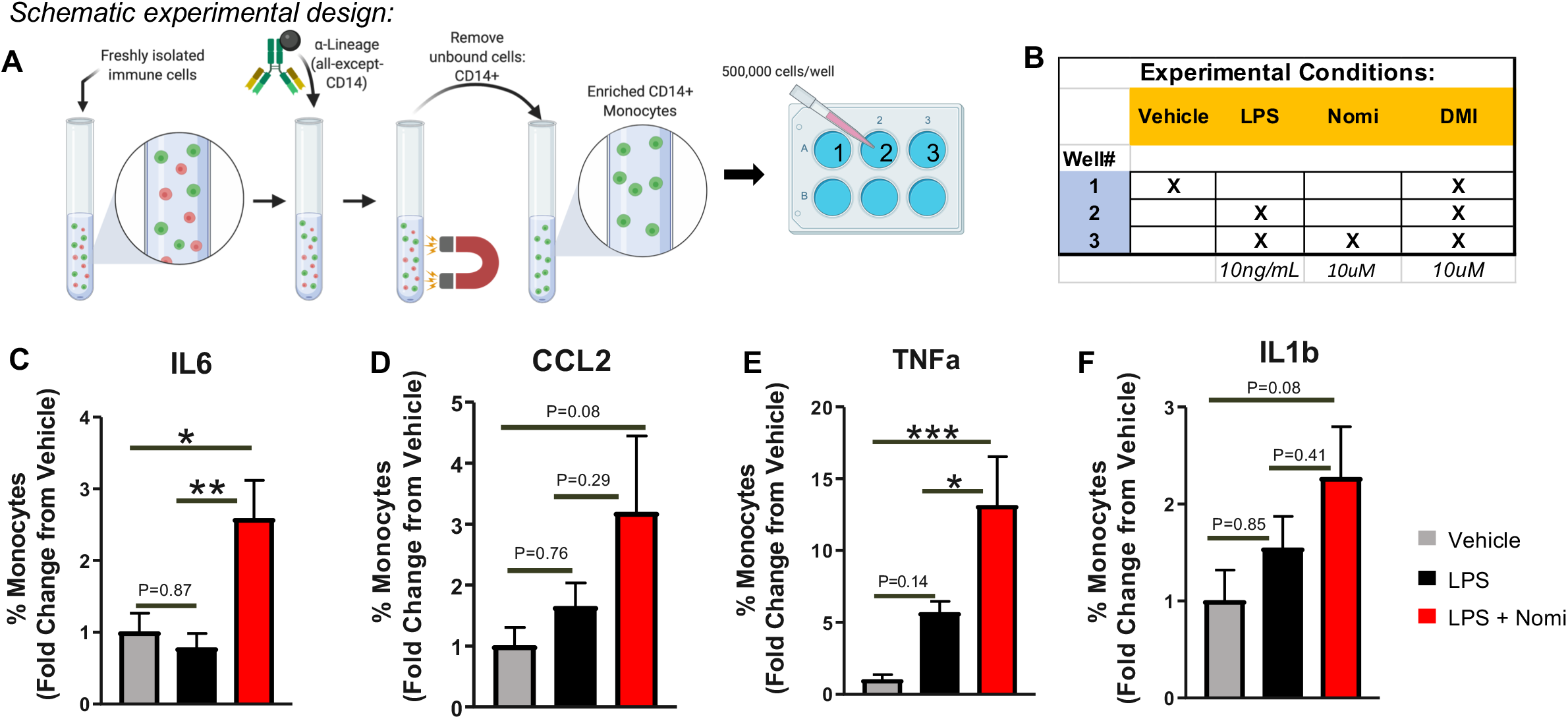
DAT inhibition enhances pro-inflammatory intracellular cytokine expression and release in acutely isolated monocytes. A) Schematic diagram describes the experimental design: Enriched CD14+ monocytes were seeded into 6-well ultra-low adherence plates at 500,000 cells/well, containing RPMI plus 7.5% heat inactivated autologous serum. B) Table outlines experimental groups: Cells were treated with vehicle, LPS (10ng/mL) or nomifensine (10uM). To isolate dopamine receptor or DAT specific effects, all experiments were performed in the presence of NET antagonist desipramine (DMI, 10uM). C-F) LPS treatment in the presence of Nomifensine significantly increases intracellular IL6 and TNFα (C & E), and both intracellular CCL2 and IL1b show similar trends (D & F, P=0.08).

**Supplemental Figure 5:**
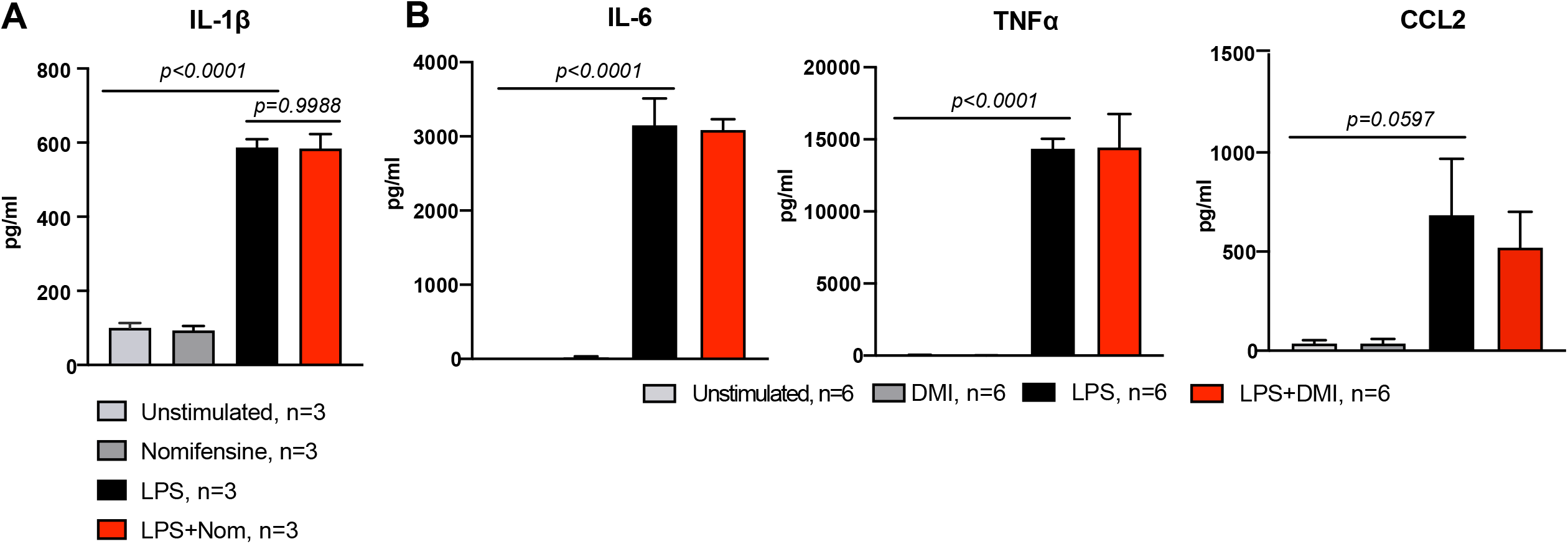
DAT inhibition has no effect on LPS-induced IL-1β, and NET inhibition does not affect LPS-induced cytokine release. (A) Lysates from macrophages treated with vehicle, nomifensine, LPS, or LPS+Nomifensine were measured for IL-1β expression via ELISA. LPS significantly increased the levels of IL-1β (p<0.0001, one-way ANOVA), and LPS+Nomifensine had no further effect (p=0.9988, one-way ANOVA); n=3 experiments/group. (B) Conditioned media from macrophages treated with vehicle (unstimulated, n=6), Desipramine (DMI, n=6), LPS (n=6), or LPS+DMI (n=6) was assayed for IL-6 (left), TNFα (middle), and CCL2 (right). LPS induced significant increase in release of pro-inflammatory cytokines and blockade of NET with DMI had no additional effect. Statistical analysis was performed using a one-way ANOVA with Tukey’s test for multiple comparisons.

**Supplemental Figure 6:**
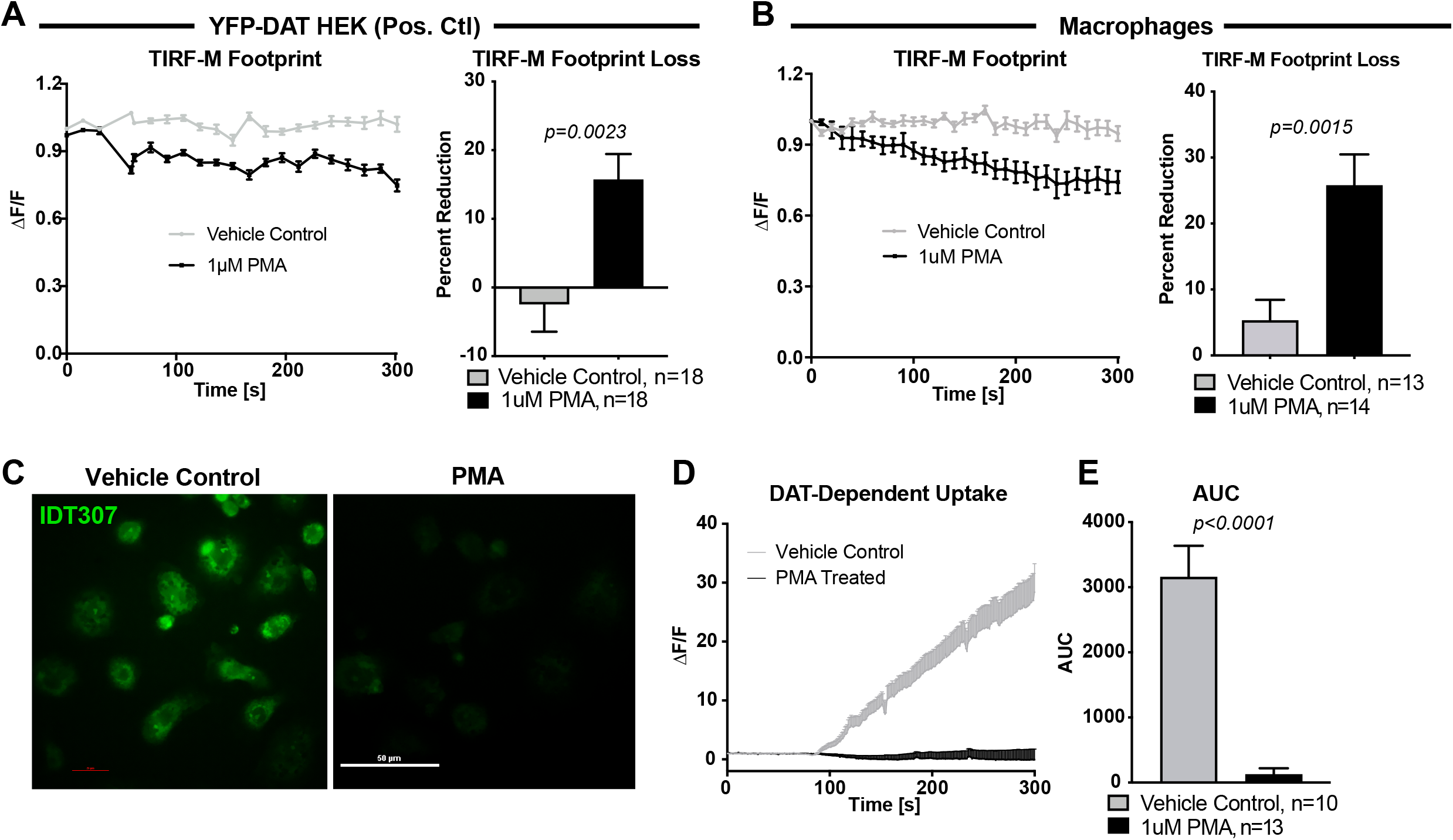
DAT on macrophages is subject to canonical regulatory mechanisms. (A) HEK cells stably transfected with YFP-DAT were incubated with JHC1-064 then treated with 1µM PMA while the JHC1-064 signal at the membrane was recorded using TIRF microscopy. Quantifying the change in JHC1-064 fluorescence over time (left) indicated a significant reduction in the JHC1-064 TIRF-M Footprint in the PMA-treated cells compared to vehicle control (p=0.0023, unpaired t-test, n=18/group). (B) Macrophages were incubated with JHC1-064 and treated with 1µM PMA while the JHC1-064 signal was recorded using TIRF microscopy. Quantifying the change in fluorescence over time (left) revealed a significant reduction in the JHC1-064 TIRF-M footprint in the PMA-treated macrophages compared to vehicle control (p=0.0015, unpaired t-test, n=13-14/group). (C) Representative images of macrophages pre-treated with either vehicle or 1µM PMA then perfused with IDT to assess uptake. (D) Blocker subtraction was performed as previously described to isolate the DAT-dependent IDT307 uptake. (E) PMA-treatment significantly reduced the magnitude of IDT307 uptake compared to vehicle control (p<0.0001, unpaired t-test, n=10-13/group).

**Supplemental Figure 7:**
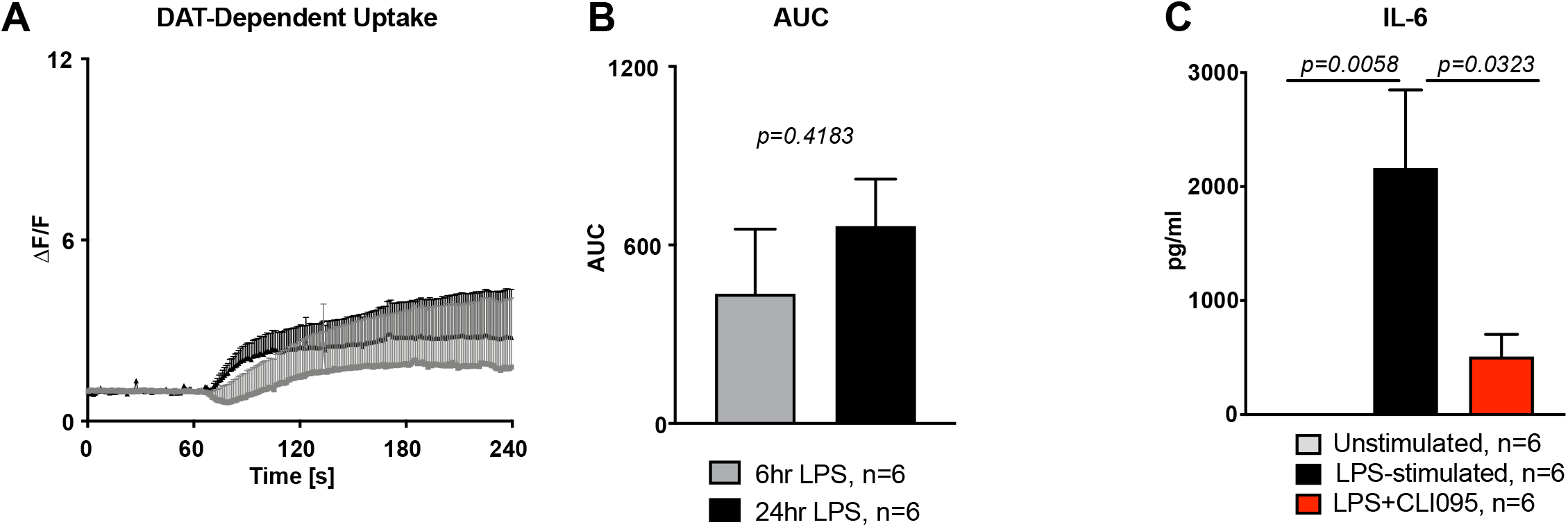
6 and 24 hour LPS-stimulated macrophages exhibit similar DAT uptake, and validation of CLI095. (A) Macrophages were treated for either 6 or 24 hours with LPS and assayed for DAT-dependent IDT307 uptake. (B) Quantification of AUC in (A) revealed that there was no difference in DAT-dependent uptake of IDT307 between the 6- and 24-hour treatments (p=0.4183, n=6, unpaired t-test). (C) Macrophages were treated with vehicle (unstimulated), LPS, or LPS + TLR4 inhibitor CLI095 and their conditioned media was assayed for IL-6 concentrations. LPS significantly increased IL6 release (p=0.0058) that was reduced with addition of CLI095 (p=0.0323, one-way ANOVA, n=6).

## Supplemental Methods

**Table S1:** Inhibitors/Fluorescent Substrates

| **Table S1: Inhibitors/Fluorescent Substrates** |  |  |  |  |  |
| --- | --- | --- | --- | --- | --- |
| **Reagent** | **Supplier** | **Cat. No.** | **Final Concentration** | **Specificity** | **Reference** |
| Nomifensine | Sigma Aldrich | N1530 | 5-20μM | IC50=7.2nM | (Rothman et al. 1993) |
| GBR | Sigma Aldrich | G9659 | 10μM | Ki=21.5nM | Sigma |
| Desipramine | Sigma Aldrich | D3900 | 1μM | Ki=4.1nM | Sigma |
| Fluoxetine | Tocris | 0927 | 1μM | Ki=0.9nM | Tocris |
| CLI-095 | Invivogen | tlrl-cl95 | 3μM | IC50=1-35nM | Li M. et al 2006 |
| IDT307 | Sigma Aldrich | SML0756 | 5μM | Km=30μM | (Blakely et al. 2011) |
| JHC1-064 | Dr. Amy Newman (NIDA) | - | 50nM | Ki=18nM | Cha et al. 2005 |
| anti-CD14 | Biolegend | 367102 | 10μg/ml | - | Kurt-Jones E.A. et al. 2000 |
| Iaxo-102 | AdipoGen | IAX-600-002-M001 | 5μM | - | (Huggins et al. 2015) |
| Sulpiride | Sigma Aldrich | S8010 | 5μM | Ki≈10nM | |
| SCH23390 | Sigma Aldrich | D054 | 5μM | Ki=0.12nM | (Faedda, Kula and Baldessarini 1989) |
| Avidin Biotin Blocking system | Biolegend | 927301 | - | - | - |

**Table S2:** Primary Antibodies/Fluorescent Dyes

| **Antibody** | **Supplier** | **Cat. No./RRID** | **Dilution** | **Application** |
| --- | --- | --- | --- | --- |
| Rat anti-DAT | Millipore | Mab369/ AB_2190413 | 1:500 | IF, WB |
| Mouse anti-TH | Millipore | T1299/ AB_477560 | 1:500 | IF |
| Rabbit anti-IBA1 | WAKO | LKF6437 | 1:500 | IF |
| CTxB-Alexa Fluor 555 | Thermo Fisher | C34776 | 1:500 | IF |
| anti-beta tubulin | Encor | MCA 1B12 | 1:40,000 | WB |
| Rabbit anti-IBA1 | Encor | RPCA-IBA1 | 1:1000,<br>1:800 | IF |
| Mouse anti-NET | MabTechnologies | NET17-1 | 1:100;<br>1:100;<br>1:1000 | IF, FC, WB |
| Mouse anti-SERT | MabTechnologies | ST51-2 | 1:2500;<br>1:100<br>1:5000 | IF, FC, WB |
| Chicken anti-MAP2 | BioLegend | Poly28225 | 1:800 | IF |
| Anti-TNF PE | BioLegend | 502908 | 1:100 | FC |
| Anti-IL6 PE-Dazzle594 | BioLegend | 501121 | 1:100 | FC |
| Anti-CCL2 APC | BioLegend | 505909 | 1:100 | FC |
| Anti-IL1β Pacific Blue | BioLegend | 511709 | 1:100 | FC |
| Fluorescent Latex Beads | Sigma Aldrich | L3280 | 1μg/ml | IF |
| MitoSox Red™ | Thermo Fisher | M36009 | 1μM | Live imaging |
| False Fluorescent Neurotransmitter 200 | Tocris | 5911 | 20μM | Live Imaging |
| Anti-Rat Dylight 405 | Novus | NBPI-72983 | 1:5000 | IF |
| Anti-Rat AlexaFluor 568 | Invitrogen | A11-077 | 1:1000 | IF |
| Anti-Rat AlexaFluor 647 | Invitrogen | A-21247 | 1:1000 | IF |
| Anti-Rat Biotin | Vector | BA9401 | 1:200 | IF |
| Anti-Mouse DyLight 405 | BioLegend | Poly24091 | 1:1000 | IF |
| Anti-Mouse AlexaFluor 647 | Invitrogen | A21235` | 1:1000 | IF |
| Anti-Rabbit AlexaFluor 568 | Invitrogen | A11011 | 1:800-1:1000 | IF |
| Anti-Chicken AlexaFluor 488+ | Invitrogen | A32931 | 1:800 | IF |
| Streptavidin AlexaFluor 647 | Invitrogen | S32357 | 1:200 | IF |

**Table S3:** Equipment used for microscopy and electrophysiology experiments

| Equipment | Experiment used | Source |
| --- | --- | --- |
| Nikon A1 laser-scanning confocal microscope | Live or Fixed cell confocal imaging | Nikon Instruments, Melville NY |
| 60x 1.49 NA Nikon Plan-Apo objective | Live or Fixed cell confocal imaging | Nikon Instruments, Melville NY |
| NIS Elements analysis software | Image analysis | Nikon Instruments, Melville NY |
| 480nm excitation laser equipped with a quadruple bandpass filter | Imaging | Chroma Technology Corp., Bellows Falls VT |
| FN-1 upright microscope | IDT307 Uptake | Warner Instruments, RC-26G |
| SpectraX Light Engine | IDT307 Uptake | Lumencor, Beaverton OR |
| Andor Zyla 4.2 PLUS camera | Imaging | Andor Technology, Belfast, UK |
| Nikon Eclipse TE 2000-U Inverted Microscope | TIRF microscopy | Nikon Instruments, Melville NY |
| 40x 1.49 NA objective | TIRF microscopy | Nikon Instruments, Melville NY |
| 40x Nikon 0.8NA Plan-APO objective | Electrophysiology and amperometry | Nikon Instruments, Melville NY |
| Amperometric carbon-fiber electrode | Amperometry | ProCFE, Dagan Corp |
| Axopatch 200B amplifier | Electrophysiology and amperometry | Molecular Devices, Sunnyvale CA |
| Digidata 1440A low-noise data acquisition system | Electrophysiology and amperometry | Molecular Devices, Sunnyvale CA |
| Axon Clampex software | Electrophysiology and amperometry | Molecular Devices, Sunnyvale CA |
| Clamp-fit software | Electrophysiology and amperometry | Sutter Instruments Co, Novato CA |
| Laser-based pipette puller, Model P-2000 | Electrophysiology and amperometry | Sutter Instruments Co, Novato CA |
| Imaging chamber | microscopy and electrophysiology | Warner Instruments, RC-26G |
| Warner VC-8 automated perfusion system | Perfusion for drug application, solution exchange | Warner Instruments, RC-26G |

